# Harnessing Community Science and Open Research-based Data to Track Distributions of Invasive Species in Japan

**DOI:** 10.1101/2024.07.30.605772

**Authors:** Shoko Sakai, Keisuke Atsumi, Takanori Genroku, Koichi Goka, Shogoro Fujiki

**Author notes:** **Corresponding author:** Shoko Sakai, Department of Geography, Hong Kong Baptist University, 15 Baptist University Rd, Kowloon Tong, Hong Kong SAR; /. **Email addresses of all authors:**.

## Abstract

At the forefront of invasive alien species (IAS) control, information gaps about the latest IAS distribution can hinder the required actions of local governments. In Japan, many prefectural governments still lack a list of invasive species despite the request stipulated in the Invasive Alien Species Management Action Plan enacted in 2015. Here, we examined to what extent open research-based data deposited by museums and herbaria (ORD) and community science data deposited by volunteers (CSD) can fill the gaps. We focused on 145 plant and 38 insect invasive species, and updated their distribution maps using ORD and CSD. We found complementarity as well as common limitations between ORD and CSD. While taxonomic biases were weaker in ORD, CSD had better prefectural coverage. In addition, some important taxa have rarely been captured by CSD or ORD. Mixed strategies of facilitating community science, supporting local museums, and taxon-specific monitoring by experts are necessary.

## INTRODUCTION

Japan is an island country that extensively depends on international trade, and imports many living organisms. As a result, the country is one of the hotspots of invasive alien species (IAS) (Mizutani and Goka 2010; Pyšek et al. 2017). About 2,200 species were listed as invasive in the country more than 20 years ago (Ecological Society Japan 2002). While it may have passed its peak (Egawa and Koyama 2023), the unceasing introduction and expansion of invasive species still threaten the biodiversity and economy of the country (Watari et al. 2021).

To tackle this issue, the Japanese government established the Invasive Alien Species Management Action Plan in 2015. This plan requests prefectural governments to list invasive species within their jurisdiction. However, as of January 2021, only 28 prefectures out of 47 had established an invasive species list (Ministry of the Environment 2021). The lack of information on IAS distribution at the prefecture level makes it difficult to determine the distribution status of IAS at the national level. While the National Institute for Environmental Studies, Japan has compiled the Invasive Species Database (ISD, https://www.nies.go.jp/biodiversity/invasive/index_en.html) including the presence or absence of >400 species in each prefecture, ISD may underestimate their distributions because it is based only on solid literature such as academic papers and local government reports.

One potential source to fill the gap is open research-based data (ORD) stored at data-sharing infrastructures. These infrastructures have been established to facilitate the digitization and mobilization of information stored at museums and herbaria (Nelson and Ellis 2018). Currently, the Global Biodiversity Information Facility (GBIF) is the largest holder among the open databases (Nelson and Ellis 2018). Since its establishment in 1999, nearly 3 billion occurrence records have been deposited in GBIF by the time of writing (May 2024).

Another approach is to leverage data generated by volunteers through community science programs. The rise of community science, also called “citizen science”, in biodiversity research has led to a surge in data collection with the establishment of online platforms such as iNaturalist in 2008 (Silvertown 2009; Bonney et al. 2014; Theobald et al. 2015; Pocock et al. 2018; Sheard et al. 2024). iNaturalist has accumulated 190 million occurrence records worldwide uploaded by 3 million observers (https://www.inaturalist.org, accessed in June 2024). In Japan, Biome, a mobile application developed by Biome Inc. in 2019, has accumulated >7 million records (Atsumi et al. 2024).

Nevertheless, multiple factors limit the information that can be extracted from massive CSD or ORD (Beck et al. 2014; Amano et al. 2016). CSD records are often concentrated in regions with high human population densities and are easily accessible (Geurts et al. 2023; but Atsumi et al. 2024). Besides, CSD is biased toward large charismatic species (e.g., mammals, birds, butterflies), and records of inconspicuous or unpopular groups (e.g., spiders) tend to be rare (Binley et al. 2023). Furthermore, CSD includes records with high uncertainties and low reliability (Aceves-Bueno et al. 2017). In contrast, the availability of ORD may depend on funding policies of the region (Yesson et al. 2007). In addition, museum collections, the main source of information for ORD, may overrepresent rare taxa since overly common species can be ignored (Guralnick and Van Cleve 2005). Therefore, it is essential to understand these characteristics when we adopt these data for invasive species monitoring.

In this study, we focused on 145 plant and 38 insect invasive species, subspecies, or varieties (we call them “species” hereafter) in ISD and evaluated the potential contributions of ORD and CSD to updating the presence and absence of the species in each prefecture in Japan. We used data from herbarium and museum specimens on GBIF as ORD, and data from iNaturalist and Biome as CSD. First, we examine to what extent ORD and CSD can identify new distributions of the target species not recognized by the ISD maps. Second, we highlight the similarity and complementarity of ORD and CSD regarding taxonomic and spatial biases. Based on the results, we discuss the advantages and limitations of ORD and CSD, and strategies to improve the monitoring of invasive species in Japan.

## MATERIALS AND METHODS

### Target species

We accessed the ISD and obtained the distribution maps of 145 vascular plant and 38 insect species in February 2023. The growth type information of the plant species was obtained from the TRY Plant Trait Database (Kattge et al. 2020). Size data for the insects was obtained from the ISD. See **Tables S1** and **S2** for further information.

### Datasets

As ORD, we downloaded occurrence records in Japan from GBIF in May 2023 and excluded records from iNaturalist. The majority of the records were ones deposited by museums and herbaria. We referred to the “state” variable of the data to identify their prefecture. If the information was unavailable, we used other locality information (the name of the city or longitude and latitude). We excluded records without prefectural or collection year information or records that were too old (<1700).

We obtained CSD from iNaturalist and Biome. In May 2023, we downloaded iNaturalist records of research grade (at least two-thirds of the community consensus on identification) through GBIF. We extracted records of the target species with valid collection year and prefecture information. Their average accuracy and precision are 96.7% and 100.0% (i.e., all records are identified into species), respectively (iNaturalist 2024).

Biome is a mobile application launched by Biome Inc. to boost citizens’ engagement in biodiversity surveys and environmental education in Japan. We used the data that had been collected by February 2023. Their species accuracy was 91.7% and 95.5% for common seed plants and insects, respectively (Atsumi et al. 2024). Before the analysis, we filtered out records deemed to be invalid based on location metadata and users’ reactions to the record (see Atsumi et al. (2024) for detailed procedures).

### Data analysis

We identified “new distributions” by comparing the ISD maps and distribution of ORD and CSD records for each species. When at least one ORD and one iNaturalist record was found in the prefecture where the species was “absent” on the ISD map, we regarded it as a new distribution. Biome records from a prefecture where neither ISD nor iNaturalist suggested the presence of the species were manually verified for the species identity and origin, and only records deemed naturalized based on photographs and locality were accepted as a new distribution. For each target species, we classified the 47 prefectures of Japan into the following five categories: 1) the prefectures where the species had already been recorded by ISD; 2) the prefectures where we newly found the presence based on both ORD and CSD, 3) based on ORD, or 4) based on CSD; and 5) the prefectures without a record. All statistical analyses in this study were conducted using the R computing environment (R Core Team 2020).

To evaluate the impact of biases in CSD and ORD, we drew rarefaction curves and compared the rate of increase in prefectures and species relative to records between CSD and ORD. We used the R package *iNEXT* (Hsieh et al. 2016) to calculate rarefaction curves with extrapolation using 1,000 bootstrap replicates. For both prefecture and species coverage, we also drew rarefaction curves using only recent ORD (year >2000) to examine the potential effects of the different time spans between ORD and CSD (see Results, **Figure 1B**).

**Figure 1.**
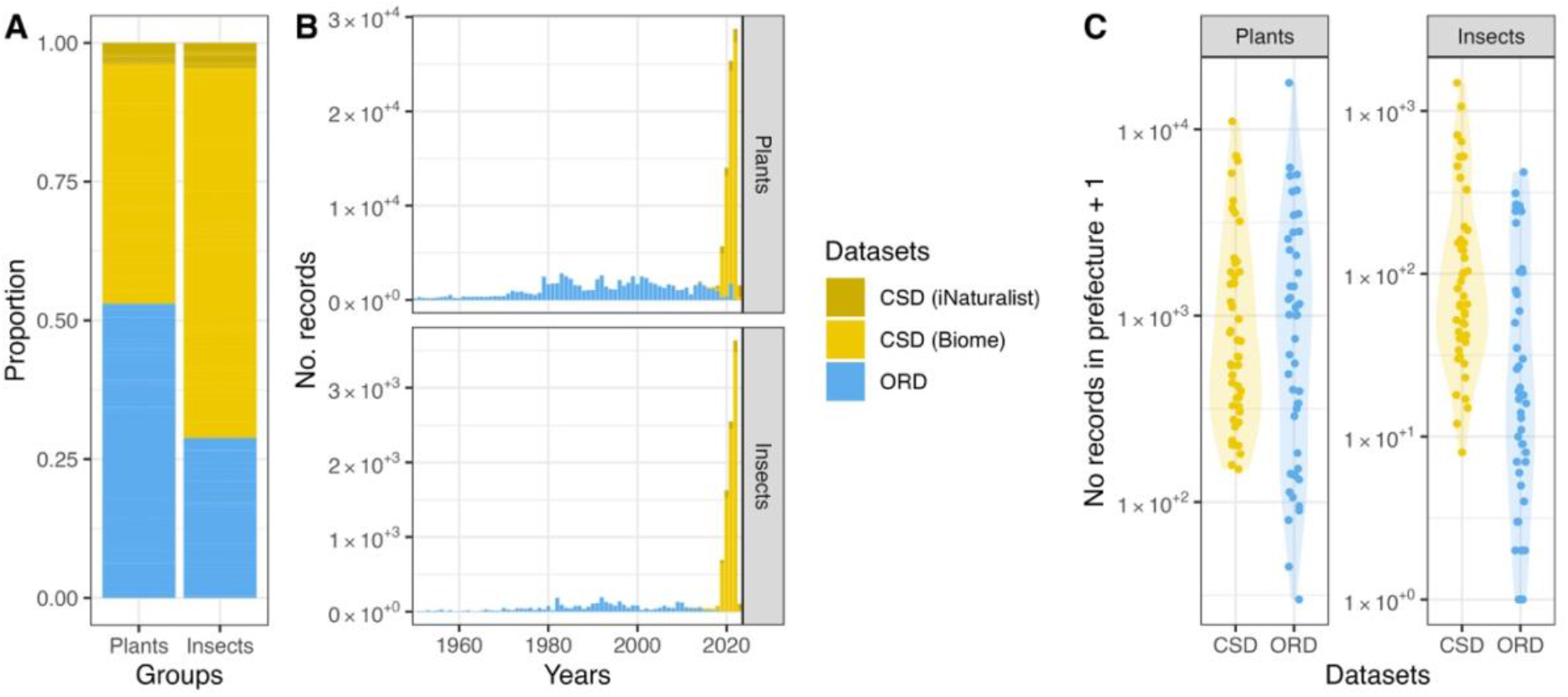
Barplots for the proportion (**A**) and temporal distribution (**B**), and a violin plot showing the distribution, of record numbers in the 47 prefectures (**C**) of community science data (CSD) and open research-based data (ORD) of the target plant and insect species. CSD is from two sources, Biome and iNaturalist. Bars before 1950 are not shown in **B**. Biome and iNaturarist data in CSD are not distinguished in **C**.

We performed Poisson regression analyses to examine whether species characters affect the record distributions among species and whether they differ between CSD and ORD. We considered datasets (CSD vs. ORD), species traits (growth types (herb vs. woody plants) for plants, log-transformed body size for insects), and their interaction as fixed terms. We included species identity as a random effect. The regression analyses and visualization were conducted using the *lme4* and *sjPlot* packages (Bates et al. 2015; Lüdecke 2021) implemented in R.

We constructed a binomial generalized linear mixed model (GLMM) to examine the relationships between the number of occurrence records and whether the occurrence of the species in the prefecture was confirmed by the validation of Biome data using the *lme4* package. The dependent variable of the model was whether the distribution was confirmed or not. The model included the Biome record numbers of the species in the prefecture and the group of the species (plant or insect) as fixed terms and the species as a random term.

## RESULTS

### Occurrence records from CSD and ORD

We obtained 157,786 occurrence records for the 145 plant species, including 83,631 of ORD and 74,155 of CSD (Biome, 68,016; iNaturalist, 6,139) (**Figure 1A**). The number for the 38 insect species was much lower, 12,352, including 3,561 of ORD and 8,791 of CSD (Biome, 8,232; iNaturalist, 559). The yearly increase in ORD was relatively stable from the 1950s to the 2000s (**Figure 1B**). In the meantime, CSD showed a rapid increase in the last several years. All dataset sources (ORD, Biome, and iNaturalist) show some degree of spatial heterogeneity (**Figure S1**). ORD and CSD record numbers were also heterogeneous among the plant and insect species as well (**Figure S2**).

### New distributions identified by ORD and CSD

This study identified potential new distributions of 106 plant invasive species among 145, and 16 insect species among 38, by combining data from CSD and ORD (**Figures 2A, S2–4**). The average number of newly recorded prefectures of these species was 4.05 for the plants and 3.94 for the insects. When only ORD was used, new distributions were found for 100 plant and 9 insect species. On the other hand, CSD indicated potentially new distributions of 78 plant and 19 insect species, while the numbers decreased to 52 and 13 after manual validation of Biome data, respectively. GLMM supported that a new distribution is more likely to be confirmed when we have more records (*z*=4.835, p=0.001 for the number of records; the effects of the group were not significant (z=–1.140, p=0.2542) (**Figure S5**). When we counted new distributions at the prefecture level, the difference between ORD and CSD was larger than that at the species level (**Figure 2B**).

**Figure 2.**
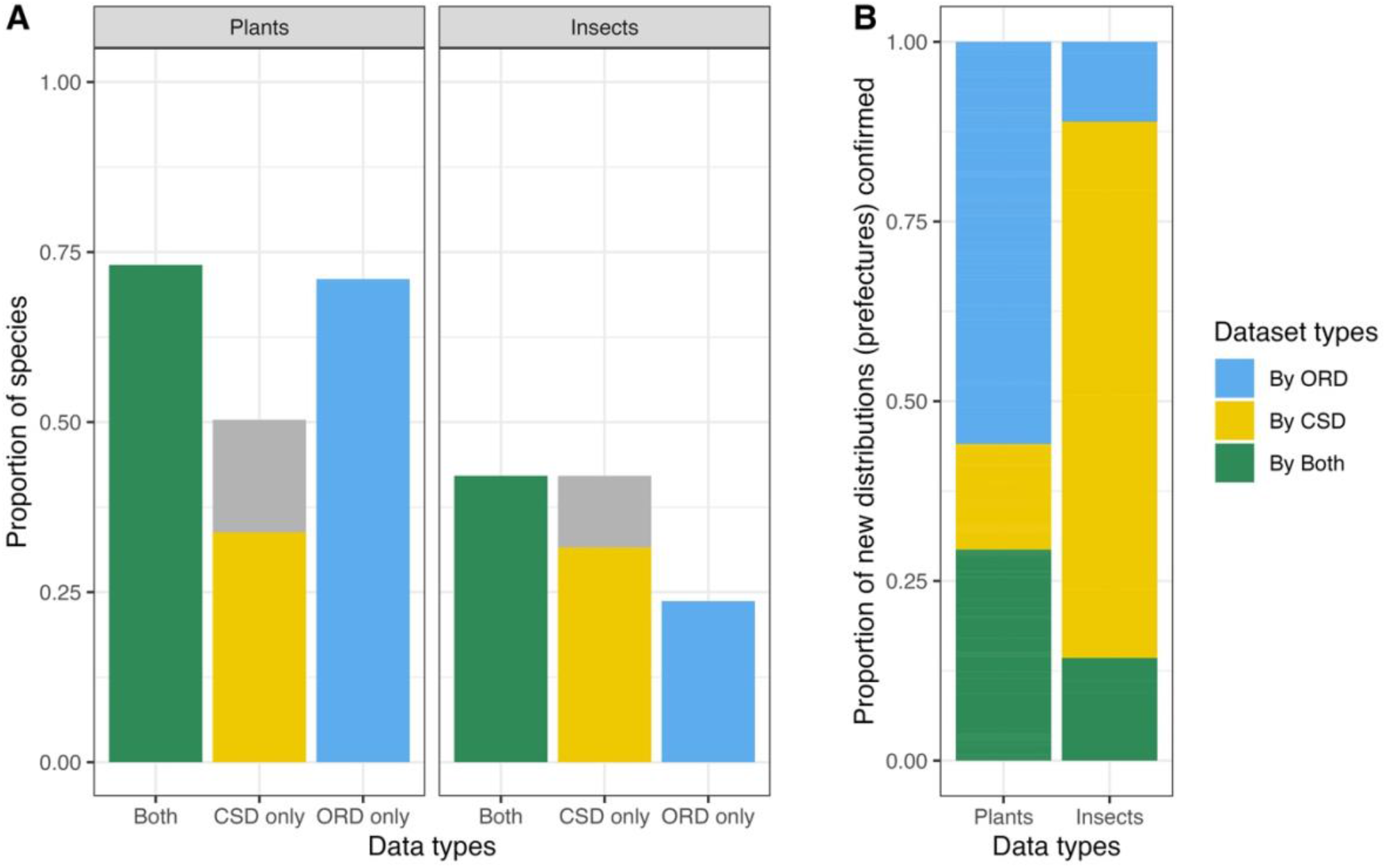
**A**. The proportion of the plant and insect species for which new distributions were confirmed. The proportions when we used both ORD and CSD (green bars), CSD only (yellow), and ORD (blue) only are shown. The portion colored with gray represents the species that had records indicating potential new distributions, but the distributions were not confirmed due to their uncertain origin or species identity (see the text). **B**. Proportions of new distributions (prefectures) confirmed by either CSD or PBD, or both CSD and PBD. In total, 429 and 63 new distributions were found for 106 plant and 16 insect species.

### Prefecture and species coverage

The rarefaction curves indicated that CSD had better prefecture coverage when the record numbers were the same both in the cases of both plants and insects (**Figure 3**). The curves for CSD were consistently higher than those for ORD, and the patterns did not change when we used only ORD in years >2000. On the other hand, for species coverage, ORD was better than CSD.

**Figure 3.**
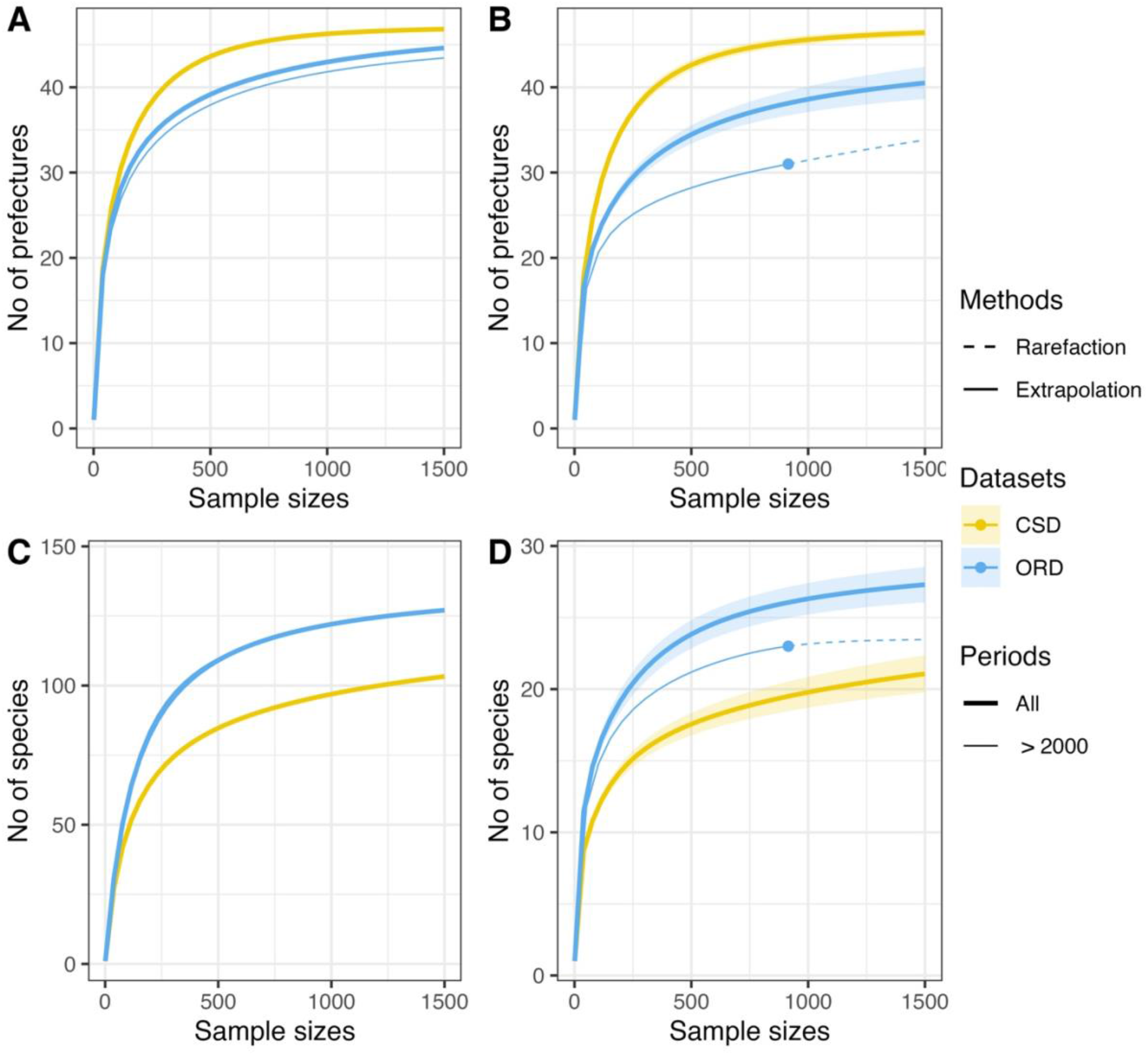
Rarefaction curves to show prefecture coverage for plants (**A**) and insects (**B**), and species coverage for plants (**C**) and insects (**D**). The rarefaction curves (solid line) are shown with extrapolation (dashed line) and 95% confidence intervals (shaded) generated with 1,000 bootstrap replicates using the R package *iNEXT*. Different colors indicate the datasets (CSD and ORD). The dot on a line in **B** and **D** indicates the end of the rarefaction. The thinner line is a curve where ORD records in the years >2000 were used.

Poisson regression analyses indicated significant differences in the record distribution between CSD and ORD. Among plants, herbs had more records than woody plants, albeit the difference was insignificant (p=0.183). Among insects, larger species had significantly more records (p<0.001). In both models, the effects of the interaction were highly significant (p<0.001) (**Table S3**). The model showed that the effect of the species traits was more pronounced in CSD than in ORD (**Figure 4**).

**Figure 4.**
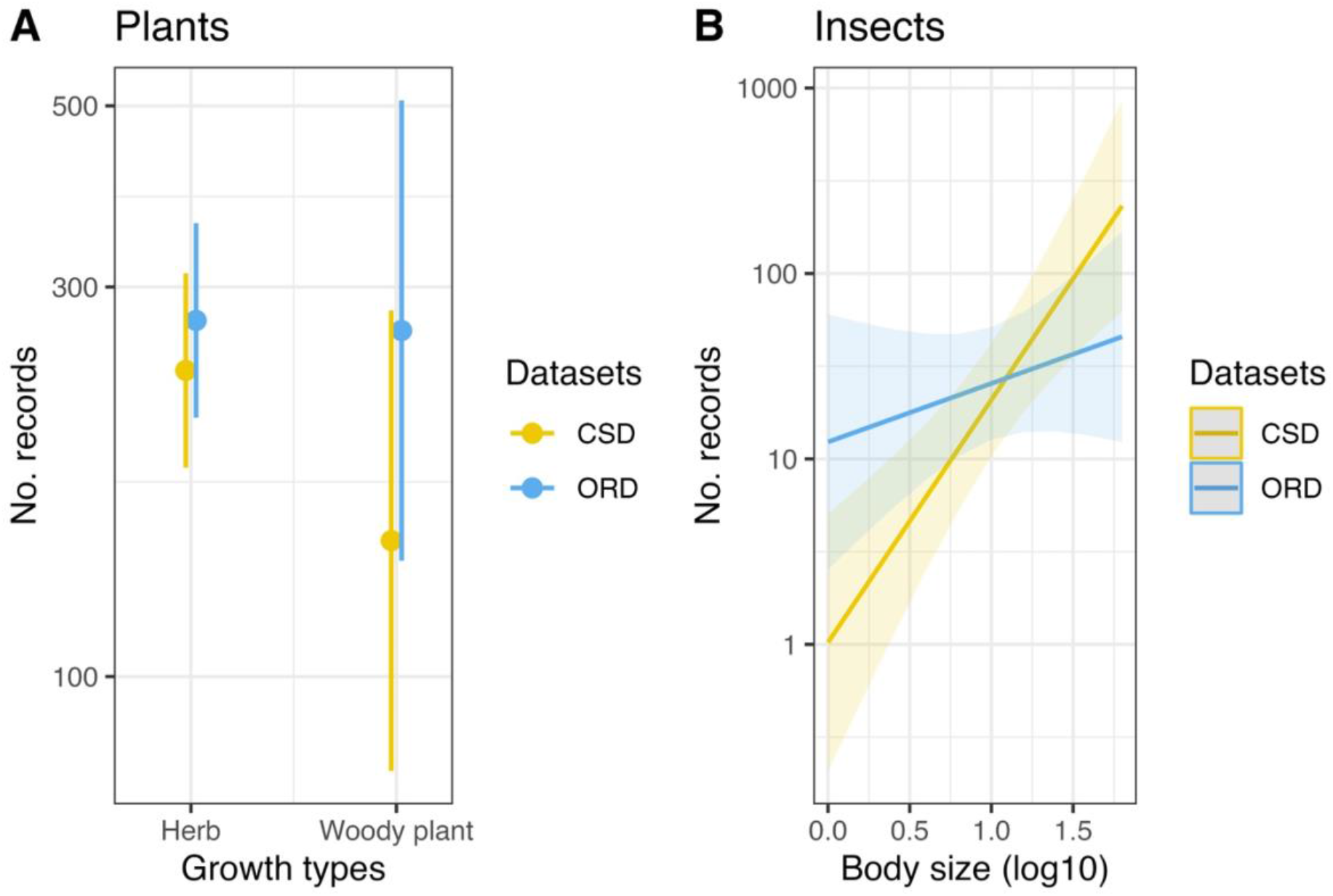
Relationships between record numbers of CSD and ORD and species trait (Growth types for plants and log-transformed body size for insects) estimated by Poisson regression analyses (**Table S3**).

## DISCUSSION

This study identified potential new distributions for 73% and 42% of the plant and insect species based on ORD and CSD, respectively (**Figures 2A, S4, S5**). Though the presence should be distinguished from the establishment of the species, the information will help the national/local governments to monitor the species and optimize additional surveys when necessary. These new distributions may include those that had been overlooked in the literature despite the species having been established for a long time, or those with which the literature had not kept up because the distribution of the species had expanded so recently. For example, many new distributions of a tree species, *Ailanthus altissima* (Simaroubaceae), recorded by both ORD and CSD, may represent the former case. In the ISD, the species is shown to be distributed across four distant regions, but the distributions captured by CSD and ORD fill the gaps between these regions (**Figure 5A**). On the other hand, a moth listed among the worst 100 invasive species, *Parasa lepida* (Lepidoptera), seems to have spread northward recently (**Figure 5B**). These new distributions were mostly identified by CSD recorded in the last several years. These results indicate that both ORD and CSD are valuable sources of information.

**Figure 5.**
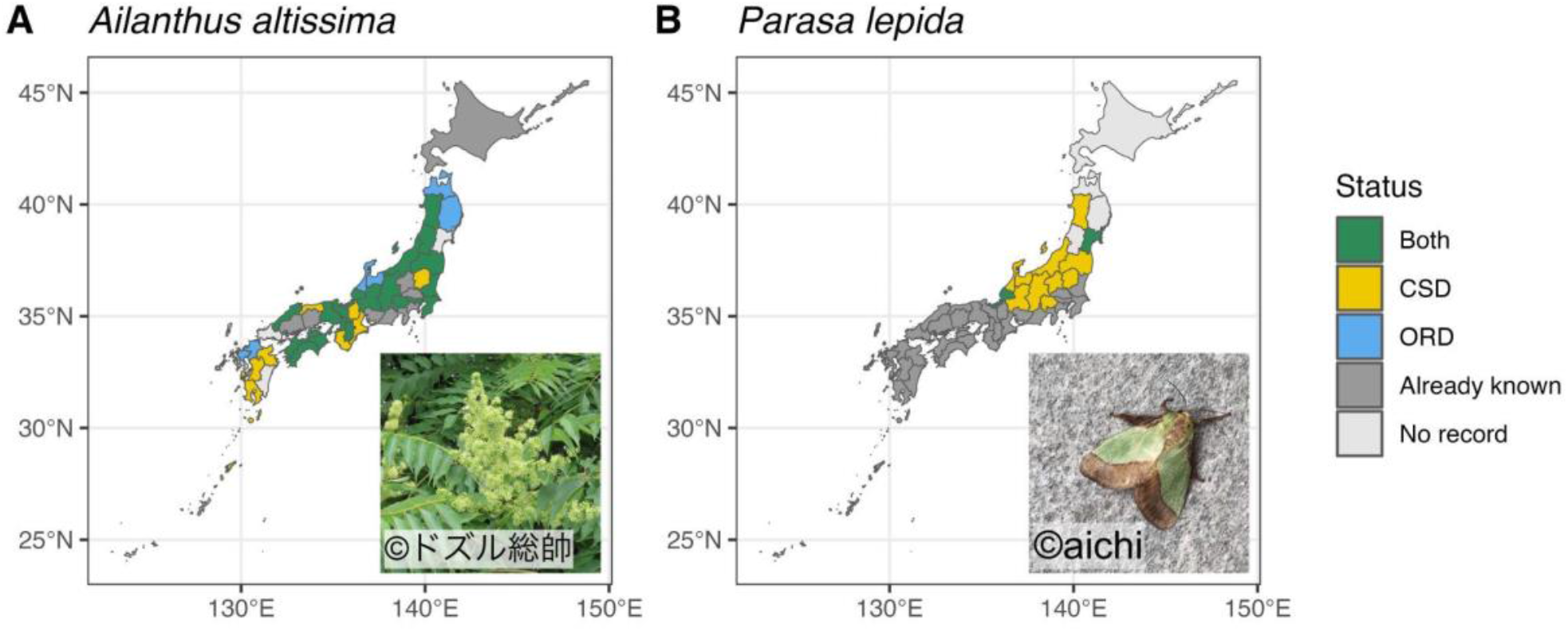
The updated distribution of a tree, *Allanthus altissima* (Simaroubaceae) (**A**) and *Parasa lepida* (Lepidoptera) (**B**). The colors of the prefectures represent the following categories: the prefecture where 1) the presence of the species had already been reported in the Invasive Species Database (ISD) (Already known), 2) the presence of species was indicated by both CSD and ORD (Both), 3) by CSD only (CSD) or 4) by ORD only, and 5) there is no data supporting the presence of the species (No record). The original shape file of the map was obtained from the Geospatial Information Authority of Japan (https://www.gsi.go.jp/top.html) (CC BY 4.0) and modified by the authors for analytical purposes. A representative photo of each species in Biome data is inserted.

We found significant differences between the relative contributions of CSD and ORD between the plants and insects. Among the plants, the new distributions were mostly identified by ORD (**Figure 2B**), whereas the total record numbers of CSD and ORD are almost equal (**Figure 1A**). In contrast, the majority (75%) of the new insect distributions were identified only by CSD, while only 11% were identified only by ORD (**Figure 2B**). Not only the difference in the record numbers but also different spatial and/or taxonomic biases of CSD and ORD in the occurrence records may affect the relative contributions.

When we compared the spatial biases based on the rarefaction curves, CSD showed greater coverage than ORD, contrary to our expectation (Geurts et al. 2023) (**Figure 2A, B**). The lower coverage of ORD may be because ORD records are considerably limited in some prefectures. Japan has as many as 723 museums with natural history specimens nationwide, but museums with sufficient resources to serve as a local hub for biodiversity monitoring are severely limited (Japanese Association of Museums 2018). Some prefectures have few records on GBIF (**Figure 1C**), possibly because they lack a hub to gather specimens and share information. In contrast, CSD is less affected by local/national policies and has few prefectures with extremely low record numbers.

Strong taxonomic biases have also been indicated for both ORD and CSD (Troudet et al. 2017; Callaghan et al. 2020) as a factor that impedes the power of detection relative to the size of the dataset. Our Poisson regression analyses suggested a strong bias toward larger insects. It is notable that CSD and ORD lack some essential insect pests that are tiny and often require well-prepared specimens and expert knowledge for identification, such as a fire ant, *Solenopsis geminata*, and a leaf-mining dipteran, *Liriomyza trifolii*. The significant interaction terms of the models for plants and insects suggested that the taxonomic biases were more pronounced in CSD (**Figure 4**). This may have caused the lower species coverage of CSD than of ORD that we observed (**Figure 3C, D**). Meanwhile, the rarefaction curve of ORD for insects could overestimate species coverage because we found that many ORD records of small insects were actually from a single collection attempt. For example, all 89 ORD records of the Argentine ant (*Linepithema humileI*) were from Tokushima on October 1st, 2010, while the species has been reported from 14 prefectures according to ISD. Unlike most vascular plants, some insect taxa require specific techniques for collecting and preparing specimens. Limited numbers of specialists and sampling opportunities may have caused these heterogeneous data distributions in insects. For better taxonomic coverage, it is important to support local museums and experts who monitor taxa rarely documented in CSD.

Frequent errors are another drawback of CSD (Aceves-Bueno et al. 2017). We strictly filtered the Biome data to record new distributions and only adopted records deemed naturalized judging from photographs (we note that records of cultivated/captive individuals can be valuable for informing the risk of naturalization (Pocock et al. 2024)). After the manual validation, the number of the plant and insect species with new distributions was reduced from 73 to 49 and 16 to 12 for the plants and insects, respectively (**Figure 2A**). Our GLMM analyses suggested that the more records we have, the more likely we will find reliable records (**Figure S5**). The progress of computer-vision-based species identifications (Saoud et al. 2020) and data filters (Bird et al. 2014; Van Eupen et al. 2021) will streamline such data validation processes. Considering the recent rapid increase of CSD, we are likely to find a way to solve this issue soon.

## CONCLUSION

We demonstrated that CSD and ORD can significantly contribute to providing information about IAS distribution. While taxonomic biases are generally weaker in ORD, CSD has better prefecture coverage, indicating the complementarity between CSD and ORD. Thus, we should make the best use of both resources. Although various research-based biodiversity data exist in Japan, only a small fraction of these data are openly available partly due to financial and logistic difficulties (Osawa 2019; Osawa et al. 2021). Currently, data from Japan accounts for only 0.5% of the GBIF occurrence data as of June 2024. In order to increase people’s involvement in invasive species monitoring through community science, communication among policy makers, researchers, community science practitioners and community scientists would be key (Maund et al. 2020; Fraisl et al. 2022). People take part in community science projects because they recognize their value for the environment and/or want to gain knowledge. Gamification is an additional feature that can boost data collection (Fraisl et al. 2022; Atsumi et al. 2024). Finally, we want to emphasize that some important taxa that require special knowledge were badly underrepresented in both CSD and ORD. This calls for taxon-specific surveys by experts organized by the national government to monitor these species.

## ACKNOWLEDGMENTS

We thank Midzuho Tatsuno, Kazumichi Morishita, and Hironori Tanaka for species identification, and Masayuki Ushio and Yuusuke Nishida for comments on an earlier version of the manuscript.

## DATA AVAILABILITY STATEMENT

Datasets and R scripts for the study are available at the Dryad Data repository via DOI: xxxxxx

