## Supplementary material for "Harnessing Community Science and Open Research-based Data to Track Distributions of Invasive Species in Japan": Table S1

### SUPPLEMENTARY TABLES

**Table S1.** The 145 target plant species with the corresponding taxon names in the ISD, Biome, and GBIF (including iNaturalist) and their growth type. For the names in the GBIF, “species” and “infraspecificEpithet” were simply combined. Among 149 plant taxa (species, varieties, and subspecies) included in ISD at the initiation of the study (February 2023), we excluded *Anthoxanthum odoratum* (Poaceae) because the taxonomic correspondence among the databases was unclear. *Lantana camara* (Verbenaceae), *Celosia argentea* (Amaranthaceae), and *Myosotis scorpioides* (Boraginaceae) were also excluded because they are frequently planted for ornamental purposes, making it difficult to distinguish between cultivated and naturalized individuals. The growth type information was obtained from the TRY Plant Trait Database (Kattge et al. 2020). The growth types of “herb,” “forb,” and “graminoid” in the original database were grouped into “herb”, and “subshrub,” “shrub,” “tree,” and “palm” into “tree.” They are indicated by 1 (herb) and 2 (woody plant). When data for the target species were unavailable, we adopted the growth habit of a congeneric species (7 species) or from the ISD website (15 species). Two of the 145 species are domestic invasive (invasive species originated from within Japan).

| Name in this study | Name in ISD | Name in GBIF | Name in Biome | Growth type |
|---|---|---|---|---|
| *Abutilon theophrasti* | *Abutilon theophrasti* | *Abutilon theophrasti* | *Abutilon theophrasti* | 1 |
| *Ageratina altissima* | *Ageratina altissima* | *Ageratina altissima* | *Ageratina altissima* | 1 |
| *Ailanthus altissima* | *Ailanthus altissima* | *Ailanthus altissima* | *Ailanthus altissima* | 2 |
| *Alternanthera denticulata* | *Alternanthera denticulata* | *Alternanthera denticulata* | *Alternanthera denticulata* | 2 |
| *Alternanthera philoxeroides* | *Alternanthera philoxeroides* | *Alternanthera philoxeroides* | *Alternanthera philoxeroides* | 1 |
| *Amaranthus retroflexus* | *Amaranthus retroflexus* | *Amaranthus retroflexus* | *Amaranthus retroflexus* | 1 |
| *Amaranthus spinosus* | *Amaranthus spinosus* | *Amaranthus spinosus* | *Amaranthus spinosus* | 1 |
| *Ambrosia artemisiifolia* | *Ambrosia artemisiifolia* | *Ambrosia artemisiifolia* | *Ambrosia artemisiifolia* | 1 |
| *Ambrosia trifida* | *Ambrosia trifida* | *Ambrosia trifida* | *Ambrosia trifida* | 2 |
| *Ammannia coccinea* | *Ammannia coccinea* | *Ammannia coccinea* | *Ammannia coccinea* | 1 |
| *Amorpha fruticosa* | *Amorpha fruticosa* | *Amorpha fruticosa* | *Amorpha fruticosa* | 2 |
| *Andropogon virginicus* | *Andropogon virginicus* | *Andropogon virginicus* | *Andropogon virginicus* | 1 |
| *Anthemis cotula* | *Anthemis cotula* | *Anthemis cotula* | *Anthemis cotula* | 1 |
| *Artemisia sieversiana* | *Artemisia sieversiana* | *Artemisia sieversiana* | *Artemisia sieversiana* | 1 |
| *Astragalus sinicus* | *Astragalus sinicus* | *Astragalus sinicus* | *Astragalus sinicus* | 1 |
| *Avena fatua* | *Avena fatua* | *Avena fatua* | *Avena fatua* | 1 |
| *Azolla cristata* | *Azolla cristata* | *Azolla cristata* | *Azolla cristata* | 1 |
| *Barbarea vulgaris* | *Barbarea vulgaris* | *Barbarea vulgaris* | *Barbarea vulgaris* | 1 |
| *Bidens frondosa* | *Bidens frondosa* | *Bidens frondosa* | *Bidens frondosa* | 1 |

| | | | | |
|---|---|---|---|---|
| *Bidens pilosa* var. *minor* | *Bidens pilosa* var. *minor* | *Bidens pilosa minor* | *Bidens pilosa* var. *minor* | 2 |
| *Bidens pilosa* var. *pilosa* | *Bidens pilosa* var. *pilosa* | *Bidens pilosa pilosa* | *Bidens pilosa* var. *pilosa* | 2 |
| *Bidens pilosa* var. *radiata* | *Bidens pilosa* var. *radiata* | *Bidens pilosa radiata* | *Bidens pilosa* var. *radiata* | 2 |
| *Bischofia javanica* | *Bischofia javanica* | *Bischofia javanica* | *Bischofia javanica* | 2 |
| *Briza maxima* | *Briza maxima* | *Briza maxima* | *Briza maxima* | 1 |
| *Bromus catharticus* | *Bromus catharticus* | *Bromus catharticus* | *Bromus catharticus* | 1 |
| *Cabomba caroliniana* | *Cabomba caroliniana* | *Cabomba caroliniana* | *Cabomba caroliniana* | 1 |
| *Casuarina equisetifolia* | *Casuarina equisetifolia* | *Casuarina equisetifolia* | *Casuarina equisetifolia* | 2 |
| *Cerastium glomeratum* | *Cerastium glomeratum* | *Cerastium glomeratum* | *Cerastium glomeratum* | 1 |
| *Chrysanthemum seticuspe* | *Chrysanthemum seticuspe* | *Chrysanthemum lavandulifolium* | *Chrysanthemum seticuspe* | 1 |
| *Cirsium arvense* | *Cirsium arvense* | *Cirsium arvense* | *Cirsium arvense* | 1 |
| *Cirsium vulgare* | *Cirsium vulgare* | *Cirsium vulgare* | *Cirsium vulgare* | 1 |
| *Conium maculatum* | *Conium maculatum* | *Conium maculatum* | *Conium maculatum* | 1 |
| *Convolvulus arvensis* | *Convolvulus arvensis* | *Convolvulus arvensis* | *Convolvulus arvensis* | 1 |
| *Corchorus olitorius* | *Corchorus olitorius* | *Corchorus olitorius* | *Corchorus olitorius* | 1 |
| *Coreopsis lanceolata* | *Coreopsis lanceolata* | *Coreopsis lanceolata* | *Coreopsis lanceolata* | 1 |
| *Cosmos bipinnatus* | *Cosmos bipinnatus* | *Cosmos bipinnatus* | *Cosmos bipinnatus* | 1 |
| *Crassocephalum crepidioides* | *Crassocephalum crepidioides* | *Crassocephalum crepidioides* | *Crassocephalum crepidioides* | 1 |
| *Cuscuta campestris* | *Cuscuta campestris* | *Cuscuta campestris* | *Cuscuta campestris* | 1 |
| *Cymbalaria muralis* | *Cymbalaria muralis* | *Cymbalaria muralis* | *Cymbalaria muralis* | 1 |
| *Cyperus congestus* | *Cyperus congestus* | *Cyperus congestus* | *Cyperus congestus* | 1 |
| *Cyperus eragrostis* | *Cyperus eragrostis* | *Cyperus eragrostis* | *Cyperus eragrostis* | 1 |
| *Cyperus esculentus* | *Cyperus esculentus* | *Cyperus esculentus* | *Cyperus esculentus* | 1 |
| *Dactylis glomerata* | *Dactylis glomerata* | *Dactylis glomerata* | *Dactylis glomerata* | 1 |
| *Datura stramonium* | *Datura stramonium* | *Datura stramonium* | *Datura stramonium* | 1 |
| *Desmodium paniculatum* | *Desmodium paniculatum* | *Desmodium paniculatum* | *Desmodium paniculatum* | 1 |
| *Drosera intermedia* | *Drosera intermedia* | *Drosera intermedia* | *Drosera intermedia* | 1 |
| *Dysphania pumilio* | *Chenopodium pumilio* | *Dysphania pumilio* | *Dysphania pumilio* | 1 |
| *Egeria densa* | *Egeria densa* | *Elodea densa* | *Egeria densa* | 1 |
| *Eichhornia crassipes* | *Eichhornia crassipes* | *Pontederia crassipes* | *Eichhornia crassipes* | 1 |
| *Elodea nuttallii* | *Elodea nuttallii* | *Elodea nuttallii* | *Elodea nuttallii* | 1 |
| *Elymus repens* | *Elytrigia repens* | *Elymus repens* | *Elytrigia repens* | 1 |
| *Eragrostis curvula* | *Eragrostis curvula* | *Eragrostis curvula* | *Eragrostis curvula* | 1 |
| *Erigeron annuus* | *Erigeron annuus* | *Erigeron annuus* | *Erigeron annuus* | 1 |
| *Erigeron canadensis* | *Conyza canadensis* | *Erigeron canadensis* | *Erigeron canadensis* | 1 |
| *Erigeron karvinskianus* | *Erigeron karvinskianus* | *Erigeron karvinskianus* | *Erigeron karvinskianus* | 1 |
| *Erigeron philadelphicus* | *Erigeron philadelphicus* | *Erigeron philadelphicus* | *Erigeron philadelphicus* | 1 |
| *Erigeron sumatrensis* | *Conyza sumatrensis* | *Erigeron sumatrensis* | *Erigeron sumatrensis* | 1 |
| *Euphorbia maculata* | *Chamaesyce maculata* | *Euphorbia maculata* | *Euphorbia maculata* | 1 |
| *Froelichia gracilis* | *Froelichia gracilis* | *Froelichia gracilis* | *Froelichia gracilis* | 1 |

| | | | | |
|---|---|---|---|---|
| *Gaillardia pulchella* | *Gaillardia pulchella* | *Gaillardia pulchella* | *Gaillardia pulchella* | 2 |
| *Galinsoga quadriradiata* | *Galinsoga quadriradiata* | *Galinsoga quadriradiata* | *Galinsoga quadriradiata* | 1 |
| *Gamochaeta coarctata* | *Gamochaeta coarctata* | *Gamochaeta coarctata* | *Gamochaeta coarctata* | 1 |
| *Gamochaeta pensylvanica* | *Gamochaeta pensylvanica* | *Gamochaeta pensylvanica* | *Gamochaeta pensylvanica* | 1 |
| *Gymnocoronis spilanthoides* | *Gymnocoronis spilanthoides* | *Gymnocoronis spilanthoides* | *Gymnocoronis spilanthoides* | 1 |
| *Helianthus tuberosus* | *Helianthus tuberosus* | *Helianthus tuberosus* | *Helianthus tuberosus* | 1 |
| *Hexapetalum teres* | *Diodia teres* | *Hexapetalum teres* | *Hexapetalum teres* | 1 |
| *Hibiscus cannabinus* | *Hibiscus cannabinus* | *Hibiscus cannabinus* | *Hibiscus cannabinus* | 1 |
| *Holcus lanatus* | *Holcus lanatus* | *Holcus lanatus* | *Holcus lanatus* | 1 |
| *Hordeum murinum* | *Hordeum murinum* | *Hordeum murinum* | *Hordeum murinum* | 1 |
| *Hydrocotyle ranunculoides* | *Hydrocotyle ranunculoides* | *Hydrocotyle ranunculoides* | *Hydrocotyle ranunculoides* | 1 |
| *Hypochaeris radicata* | *Hypochaeris radicata* | *Hypochaeris radicata* | *Hypochaeris radicata* | 1 |
| *Ipomoea coccinea* | *Ipomoea coccinea* | *Ipomoea coccinea* | *Ipomoea coccinea* | 1 |
| *Ipomoea hederacea* | *Ipomoea hederacea* | *Ipomoea hederacea* | *Ipomoea hederacea* | 1 |
| *Iris pseudacorus* | *Iris pseudacorus* | *Iris pseudacorus* | *Iris pseudacorus* | 1 |
| *Lamium purpureum* | *Lamium purpureum* | *Lamium purpureum* | *Lamium purpureum* | 1 |
| *Lepidium didymum* | *Coronopus didymus* | *Lepidium didymum* | *Lepidium didymum* | 1 |
| *Leucaena leucocephala* | *Leucaena leucocephala* | *Leucaena leucocephala* | *Leucaena leucocephala* | 2 |
| *Ligustrum lucidum* | *Ligustrum lucidum* | *Ligustrum lucidum* | *Ligustrum lucidum* | 2 |
| *Lilium formosanum* | *Lilium formosanum* | *Lilium formosanum* | *Lilium formosanum* | 1 |
| *Lindernia dubia* | *Lindernia dubia* | *Lindernia dubia* | *Lindernia dubia* | 1 |
| *Lolium arundinaceum* | *Festuca arundinacea* | *Lolium arundinaceum* | *Lolium arundinaceum* | 1 |
| *Lolium multiflorum* | *Lolium multiflorum* | *Lolium multiflorum* | *Lolium multiflorum* | 1 |
| *Lotus corniculatus* var. *corniculatus* | *Lotus corniculatus* var. *corniculatus* | *Lotus corniculatus corniculatus* | *Lotus corniculatus* var. *corniculatus* | 1 |
| *Ludwigia grandiflora* | *Ludwigia grandiflora* | *Ludwigia hexapetala, Ludwigia grandiflora* | *Ludwigia grandiflora* | 1 |
| *Medicago lupulina* | *Medicago lupulina* | *Medicago lupulina* | *Medicago lupulina* | 1 |
| *Mikania micrantha* | *Mikania micrantha* | *Mikania micrantha* | *Mikania micrantha* | 1 |
| *Mirabilis jalapa* | *Mirabilis jalapa* | *Mirabilis jalapa* | *Mirabilis jalapa* | 1 |
| *Morus australis* | *Morus australis* | *Morus australis* | *Morus australis* | 2 |
| *Myriophyllum aquaticum* | *Myriophyllum aquaticum* | *Myriophyllum aquaticum* | *Myriophyllum aquaticum* | 1 |
| *Nasturtium officinale* | *Nasturtium officinale* | *Nasturtium officinale* | *Nasturtium officinale* | 1 |
| *Nothoscordum gracile* | *Nothoscordum gracile* | *Nothoscordum gracile* | *Nothoscordum gracile* | 1 |
| *Nuttallanthus canadensis* | *Nuttallanthus canadensis* | *Nuttallanthus canadensis* | *Nuttallanthus canadensis* | 1 |
| *Oenothera biennis* | *Oenothera biennis* | *Oenothera biennis* | *Oenothera biennis* | 1 |
| *Oenothera laciniata* | *Oenothera laciniata* | *Oenothera laciniata* | *Oenothera laciniata* | 1 |
| *Oenothera rosea* | *Oenothera rosea* | *Oenothera rosea* | *Oenothera rosea* | 1 |
| *Orobanche minor* | *Orobanche minor* | *Orobanche minor* | *Orobanche minor* | 1 |
| *Orychophragmus violaceus* | *Orychophragmus violaceus* | *Orychophragmus violaceus* | *Orychophragmus violaceus* | 1 |
| *Oxalis debilis* subsp. *corymbosa* | *Oxalis corymbosa* | *Oxalis debilis corymbosa* | *Oxalis debilis* subsp. *corymbosa* | 2 |

| | | | | |
|---|---|---|---|---|
| *Oxalis pes-caprae* | *Oxalis pes-caprae* | *Oxalis pes-caprae* | *Oxalis pes-caprae* | 1 |
| *Papaver dubium* | *Papaver dubium* | *Papaver dubium* | *Papaver dubium* | 1 |
| *Paspalum distichum* | *Paspalum distichum* | *Paspalum distichum* | *Paspalum distichum* | 1 |
| *Petrorhagia dubia* | *Petrorhagia nanteulii* | *Petrorhagia dubia* | *Petrorhagia dubia* | 1 |
| *Phleum pratense* | *Phleum pratense* | *Phleum pratense* | *Phleum pratense* | 1 |
| *Phyllostachys edulis* | *Phyllostachys edulis* | *Phyllostachys edulis* | *Phyllostachys edulis* | 1 |
| *Physalis angulata* | *Physalis angulata* | *Physalis angulata* | *Physalis angulata* | 1 |
| *Phytolacca americana* | *Phytolacca americana* | *Phytolacca americana* | *Phytolacca americana* | 1 |
| *Pinus luchuensis* | *Pinus luchuensis* | *Pinus luchuensis* | *Pinus luchuensis* | 2 |
| *Pistia stratiotes* | *Pistia stratiotes* | *Pistia stratiotes* | *Pistia stratiotes* | 1 |
| *Plantago lanceolata* | *Plantago lanceolata* | *Plantago lanceolata* | *Plantago lanceolata* | 1 |
| *Poa pratensis* | *Poa pratensis* | *Poa humilis, Poa pratensis* | *Poa pratensis* | 1 |
| *Robinia pseudoacacia* | *Robinia pseudoacacia* | *Robinia pseudoacacia* | *Robinia pseudoacacia* | 2 |
| *Rudbeckia laciniata* | *Rudbeckia laciniata* | *Rudbeckia laciniata* | *Rudbeckia laciniata* | 2 |
| *Rumex obtusifolius* | *Rumex obtusifolius* | *Rumex obtusifolius* | *Rumex obtusifolius* | 1 |
| *Salpichroa origanifolia* | *Salpichroa origanifolia* | *Salpichroa origanifolia* | *Salpichroa origanifolia* | 1 |
| *Salvinia molesta* | *Salvinia molesta* | *Salvinia molesta* | *Salvinia molesta* | 1 |
| *Senecio madagascariensis* | *Senecio madagascariensis* | *Senecio madagascariensis* | *Senecio madagascariensis* | 1 |
| *Senecio vulgaris* | *Senecio vulgaris* | *Senecio vulgaris* | *Senecio vulgaris* | 1 |
| *Sicyos angulatus* | *Sicyos angulatus* | *Sicyos angulatus* | *Sicyos angulatus* | 1 |
| *Solanum carolinense* | *Solanum carolinense* | *Solanum carolinense* | *Solanum carolinense* | 2 |
| *Solidago altissima* | *Solidago altissima* | *Solidago altissima* | *Solidago altissima* | 1 |
| *Solidago gigantea* subsp. *serotina* | *Solidago gigantea* subsp. *serotina* | *Solidago gigantea serotina* | *Solidago gigantea* subsp. *serotina* | 2 |
| *Sonchus asper* | *Sonchus asper* | *Sonchus asper* | *Sonchus asper* | 1 |
| *Sorghum propinquum* | *Sorghum halepense* | *Sorghum propinquum* | *Sorghum propinquum* | 1 |
| *Sphagneticola trilobata* | *Sphagneticola trilobata* | *Sphagneticola trilobata* | *Sphagneticola trilobata* | 1 |
| *Sporobolus alterniflorus* | *Spartina alterniflora* | *Sporobolus alterniflorus* | *Spartina alterniflora* | 1 |
| *Symphyotrichum subulatum* | *Aster subulatus* | *Symphyotrichum squamatum, Symphyotrichum subulatum* | *Symphyotrichum subulatum* | 1 |
| *Taraxacum officinale* | *Taraxacum officinale* | *Taraxacum officinale* | *Taraxacum officinale* | 1 |
| *Trachycarpus fortunei* | *Trachycarpus fortunei* | *Trachycarpus fortunei* | *Trachycarpus fortunei* | 2 |
| *Tradescantia fluminensis* | *Tradescantia flumiensis* | *Tradescantia fluminensis* | *Tradescantia fluminensis* | 1 |
| *Trifolium repens* | *Trifolium repens* | *Trifolium repens* | *Trifolium repens* | 1 |
| *Triodanis perfoliata* | *Triodanis perfoliata* | *Triodanis perfoliata* | *Triodanis perfoliata* | 1 |
| *Tripleurospermum maritimum* subsp. *inodorum* | *Tripleurospermum maritimum* subsp. *inodorum* | *Tripleurospermum inodorum* | *Tripleurospermum maritimum* subsp. *inodorum* | 1 |
| *Ulex europaeus* | *Ulex europaeus* | *Ulex europaeus* | *Ulex europaeus* | 2 |
| *Utricularia inflata* | *Utricularia inflata* | *Utricularia inflata* | *Utricularia inflata* | 1 |
| *Valerianella locusta* | *Valerianella locusta* | *Valerianella locusta* | *Valerianella locusta* | 1 |
| *Verbascum blattaria* | *Verbascum blattaria* | *Verbascum blattaria* | *Verbascum blattaria* | 1 |

| | | | | |
|---|---|---|---|---|
| *Verbascum thapsus* | *Verbascum thapsus* | *Verbascum thapsus* | *Verbascum thapsus* | 1 |
| *Verbena bonariensis* | *Verbena bonariensis* | *Verbena bonariensis* | *Verbena bonariensis* | 1 |
| *Verbena brasiliensis* | *Verbena brasiliensis* | *Verbena brasiliensis* | *Verbena brasiliensis* | 2 |
| *Verbesina alternifolia* | *Verbesina alternifolia* | *Verbesina alternifolia* | *Verbesina alternifolia* | 1 |
| *Veronica anagallis-aquatica* | *Veronica anagallis-aquatica* | *Veronica anagallis-aquatica* | *Veronica anagallis-aquatica* | 1 |
| *Veronica arvensis* | *Veronica arvensis* | *Veronica arvensis* | *Veronica arvensis* | 1 |
| *Veronica persica* | *Veronica persica* | *Veronica persica* | *Veronica persica* | 1 |
| *Wolffia globosa* | *Wolffia globosa* | *Wolffia globosa* | *Wolffia globosa* | 1 |
| *Xanthium occidentale* | *Xanthium occidentale* | *Xanthium orientale* | *Xanthium occidentale* | 1 |
