## Supplementary material for "Harnessing Community Science and Open Research-based Data to Track Distributions of Invasive Species in Japan": Table S2

**Table S2.** The 38 target insect species with the corresponding taxon names in ISD, Biome, and GBIF (including iNaturalist), and their body size. For the names in the GBIF, “species” and “infraspecificEpithet” were simply combined. Among 41 insect species included in ISD at the initiation of the study (February 2023), *Psacothea hilaris* subspp. (Coleoptera) were excluded because of their uncertain identity. *Ropalidia marginata* (Hymenoptera), which has very limited known distribution (Ioto Island), and *Bombus terrestris* (Hymenoptera), which is often maintained in greenhouses to facilitate pollination, were excluded from the analyses. The size of the target insect species was obtained from the ISD. The body length was adopted for most species, while the length of the forewing or half of the wing span was adopted for Lepidopteran insects. When the size was indicated as a range, or when different sizes were provided for different sexes, castes, or seasons, their maximum value, except the size for the queen in social insects, was adopted. Three of the 38 species are domestic invasive.

| Name in this study | Name in ISD | Name in GBIF | Name in Biome | Body size |
|---|---|---|---|---|
| *Agriosphodrus dohrni* | *Agriosphodrus dohrni* | *Agriosphodrus dohrni* | *Agriosphodrus dohrni* | 24 |
| *Anoplophora glabripennis* | *Anoplophora glabripennis* | *Anoplophora glabripennis* | *Anoplophora glabripennis* | 35 |
| *Aromia bungii* | *Aromia bungii* | *Aromia bungii* | *Aromia bungii* | 38 |
| *Bactrocera cucurbitae* | *Bactrocera cucurbitae* | *Bactrocera cucurbitae* | *Bactrocera cucurbitae* | 8 |
| *Bactrocera dorsalis* | *Bactrocera dorsalis* | *Bactrocera dorsalis* | *Bactrocera dorsalis* | 8 |
| *Bemisia argentifolii* | *Bemisia argentifolii* | *Bemisia argentifolii* | *Bemisia argentifolii* | 1 |
| *Blattella germanica* | *Blattella germanica* | *Blattella germanica* | *Blattella germanica* | 15 |
| *Caverelius saccharivorus* | *Caverelius saccharivorus* | *Caverelius saccharivorus* | *Caverelius saccharivorus* | 8 |
| *Coptotermes formosanus* | *Coptotermes formosanus* | *Coptotermes formosanus* | *Coptotermes formosanus* | 8.5 |
| *Cylas formicarius* | *Cylas formicarius* | *Cylas formicarius* | *Cylas formicarius* | 7 |
| *Delta pyriforme* | *Delta pyriforme* | *Delta pyriforme* | *Delta pyriforme* | 28 |
| *Dryocosmus kuriphilus* | *Dryocosmus kuriphilus* | *Dryocosmus kuriphilus* | *Dryocosmus kuriphilus* | 3 |
| *Epilachna varivestis* | *Epilachna varivestis* | *Epilachna varivestis* | *Epilachna varivestis* | 8.5 |
| *Euscepes batatae* | *Euscepes postfasciatus* | *Euscepes postfasciatus* | *Euscepes batatae* | 3.6 |
| *Frankliniella occidentalis* | *Frankliniella occidentalis* | *Frankliniella occidentalis* | *Frankliniella occidentalis* | 1.7 |
| *Hestina assimilis* | *Hestina assimilis* | *Hestina assimilis* | *Hestina assimilis* | 53 |
| *Hypera postica* | *Hypera postica* | *Hypera postica* | *Hypera postica* | 6.5 |
| *Hyphantria cunea* | *Hyphantria cunea* | *Hyphantria cunea* | *Hyphantria cunea* | 21 |
| *Lepisiota frauenfeldi* | *Lepisiota frauenfeldi* | *Lepisiota frauenfeldi* | *Lepisiota frauenfeldi* | 4 |
| *Linepithema humile* | *Linepithema humile* | *Linepithema humile* | *Linepithema humile* | 2.5 |
| *Liriomyza trifolii* | *Liriomyza trifolii* | *Liriomyza trifolii* | *Liriomyza trifolii* | 2 |
| *Lissorhoptrus oryzophilus* | *Lissorhoptrus oryzophilus* | *Lissorhoptrus oryzophilus* | *Lissorhoptrus oryzophilus* | 3.5 |
| *Neil Somaiya rufella* | *Neil Somaiya rufella* | *Nealsomyia rufella* | *Nealsomyia rufella* | 5 |
| *Opisthoplatia orientalis* | *Opisthoplatia orientalis* | *Opisthoplatia orientalis* | *Opisthoplatia orientalis* | 33 |
| *Paraglenea fortunei* | *Paraglenea fortunei* | *Paraglenea fortunei* | *Paraglenea fortunei* | 20 |

| | | | | |
|---|---|---|---|---|
| *Parasa lepida* | *Parasa lepida* | *Parasa lepida* | *Parasa lepida* | 20 |
| *Protaetia orientalis sakaii* | *Protaetia orientalis sakaii* | *Protaetia orientalis sakaii* | *Protaetia orientalis sakaii* | 25 |
| *Protaetia pryeri* subsp. *Oschimana* | *Protaetia pryeri oschimana* | *Protaetia pryeri oschimana* | *Protaetia pryeri* subsp. *oschimana* | 28 |
| *Psacothea hilaris* subsp. *maculata* | *Psacothea hilaris maculata* | *Psacothea hilaris maculata* | *Psacothea hilaris* subsp. *maculata* | 30 |
| *Rhabdoscelus obscurus* | *Rhabdoscelus obscurus* | *Rhabdoscelus obscurus* | *Rhabdoscelus obscurus* | 13 |
| *Rhynchophorus ferrugineus* | *Rhynchophorus ferrugineus* | *Rhynchophorus ferrugineus* | *Rhynchophorus ferrugineus* | 35 |
| *Sericinus montela* | *Sericinus montela* subspp. | *Sericinus montela* | *Sericinus montela* | 38 |
| *Solenopsis geminata* | *Solenopsis geminata* | *Solenopsis geminata* | *Solenopsis geminata* | 8 |
| *Thrips palmi* | *Thrips palmi* | *Thrips palmi* | *Thrips palmi* | 1.1 |
| *Trialeurodes vaporariorum* | *Trialeurodes vaporariorum* | *Trialeurodes vaporariorum* | *Trialeurodes vaporariorum* | 1.2 |
| *Unaspis yanonensis* | *Unaspis yanonensis* | *Unaspis yanonensis* | *Unaspis yanonensis* | 5.5 |
| *Vespa velutina* | *Vespa velutina* | *Vespa velutina* | *Vespa velutina* | 24 |
| *Xylocopa tranquebarorum* | *Xylocopa tranquebarorum* | *Xylocopa tranquebarorum* | *Xylocopa tranquebarorum* subsp. *tranquebatorum* | 20 |
