## Supplementary material for "Harnessing Community Science and Open Research-based Data to Track Distributions of Invasive Species in Japan": Table S3

**Table S3.** The results of the Poisson regression analysis. For categorical variables, the reference level is shown in parenthesis.

Plants

| Fixed effects | Estimate | Std. Error | z value | p value | |
|---|---|---|---|---|---|
| Intercept | 5.468 | 0.1400 | 39.07 | < 0.001 | *** |
| Datasets (ORD) | 0.141 | 0.0054 | 25.88 | < 0.001 | *** |
| Growth types (Herb) | -0.480 | 0.3597 | -1.33 | 0.182 | |
| Datasets:Growth type | 0.452 | 0.0168 | 26.84 | < 0.001 | *** |

Insects

| Fixed effects | Estimate | Std. Error | z value | p value | |
|---|---|---|---|---|---|
| Intercept | 0.025 | 0.8138 | 0.030 | 0.976 | |
| Datasets (ORD) | 2.490 | 0.1112 | 22.388 | < 0.001 | *** |
| Body size | 3.012 | 0.7197 | 4.185 | < 0.001 | *** |
| Datasets:Body size | -2.286 | 0.0881 | -28.061 | < 0.001 | *** |
