## Supplementary material for "Harnessing Community Science and Open Research-based Data to Track Distributions of Invasive Species in Japan": Figure S1

### SUPPLEMENTARY FIGURES

**Figure S1.** Spatial distribution of ORD, Biome, and iNaturalist occurrence records. The color on the map indicates the proportion of the records in each prefecture. The original shape file of the map was downloaded from the Geospatial Information Authority of Japan (GSI) (https://www.gsi.go.jp/top.html) (CC BY 4.0) and modified by the authors for analytical purposes.

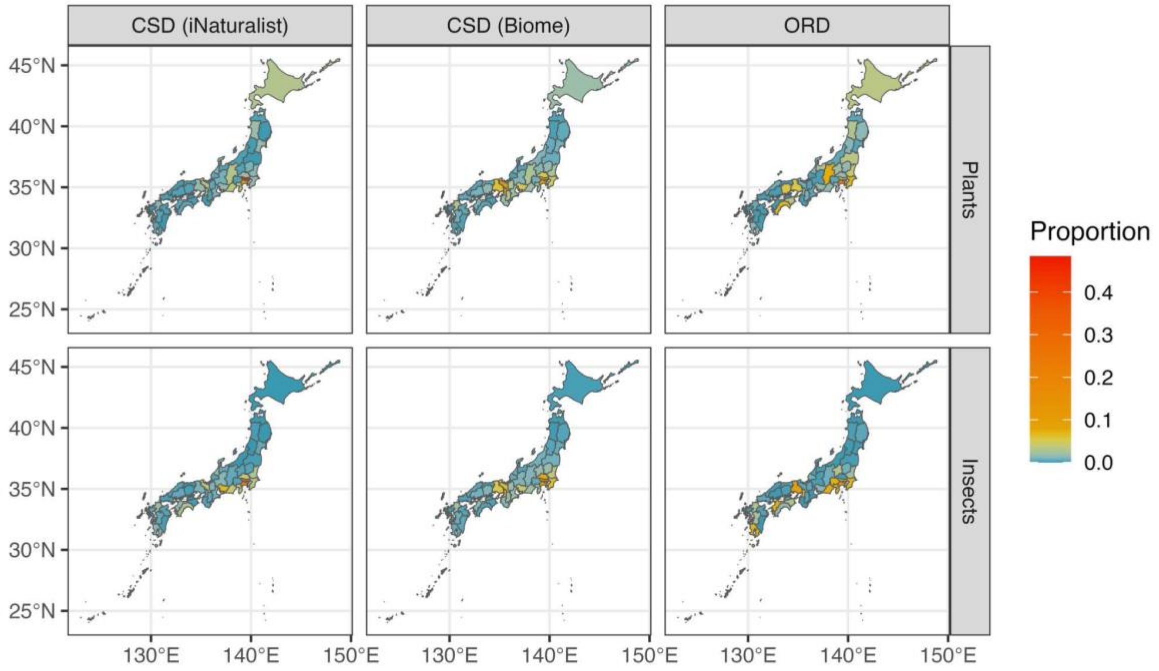
