## Supplementary material for "Harnessing Community Science and Open Research-based Data to Track Distributions of Invasive Species in Japan": Figure S2

**Figure S2.** The number of records of the target plants (**A**) and insects (**C**) in ORD and CSD, and updated distributions of the plants (**B**) and insects (**D**). In **B** and **D**, prefectures where the presence of the species had been recorded in ISD (Recorded in ISD), where presence was indicated by both ORD and CSD (ORD and CSD), by CSD only (CSD), and by ORD only (ORD), and where the presence was not supported by any data source (No record) are distinguished with different colors. Species are ordered by the number of records in CSD.

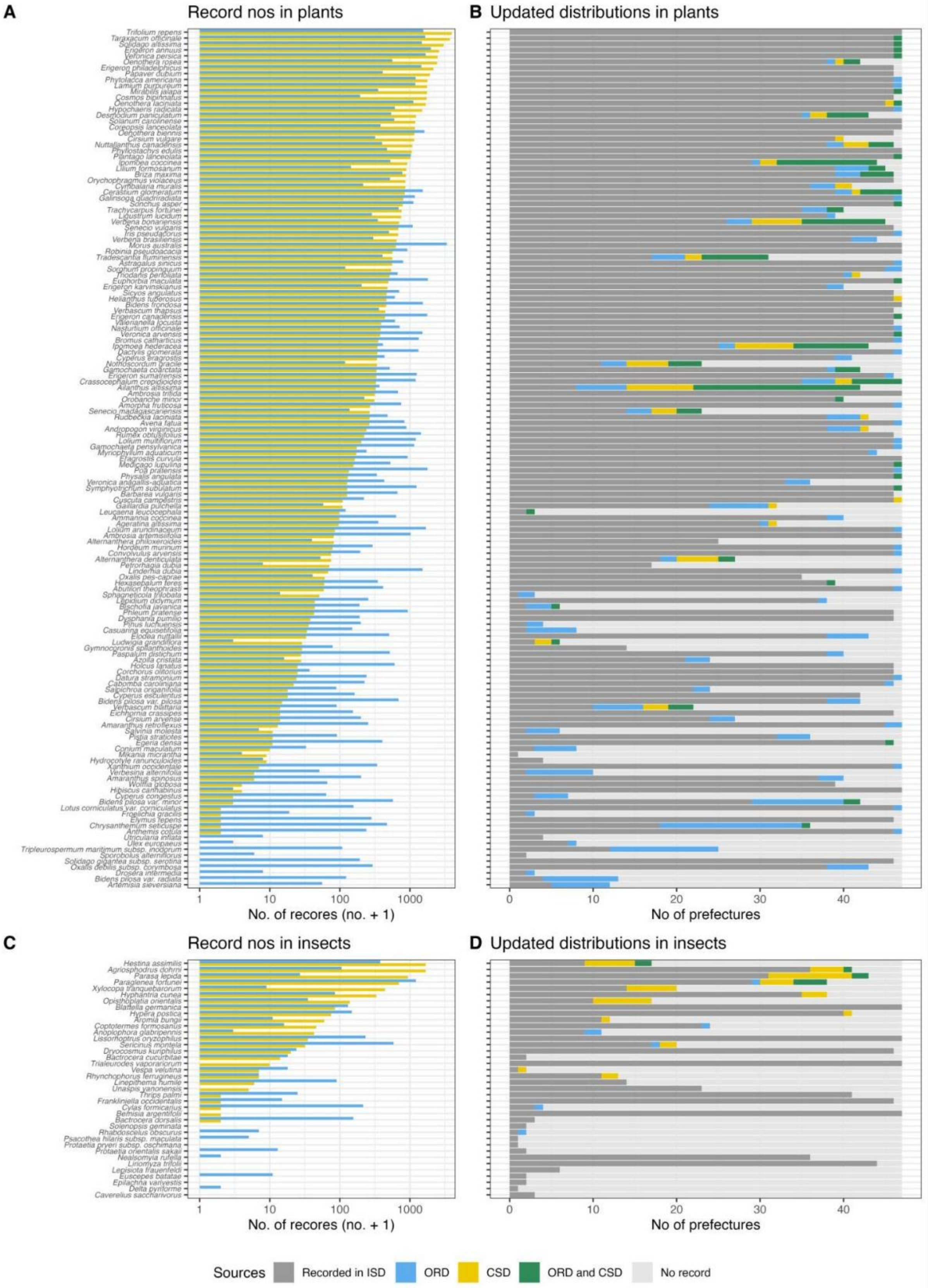
A
Record nos in plants
No. of recores (no. + 1)
1
10
100
1000
B
Updated distributions in plants
No of prefectures
0
10
20
30
40
C
Record nos in insects
No. of recores (no. + 1)
D
Updated distributions in insects
No of prefectures
Sources
Recorded in ISD
ORD
CSD
ORD and CSD
No record
