## Supplementary material for "Harnessing Community Science and Open Research-based Data to Track Distributions of Invasive Species in Japan": Figure S3

**Figure S3.** Updated maps of the 106 invasive plant species. The species in which we did not find new distributions were excluded. The colors of the prefectures represent the following categories: the prefecture where 1) the presence of the species had already been reported in the Invasive Species Database (ISD) (Already known), 2) the presence of species was indicated by both CSD and ORD (Both), 3) by CSD only (CSD) or 4) by ORD only, and 5) there is no data supporting the presence of the species (No record). The original shape file of the map was obtained from the Geospatial Information Authority of Japan (https://www.gsi.go.jp/top.html) and modified by the authors for analytical purposes.

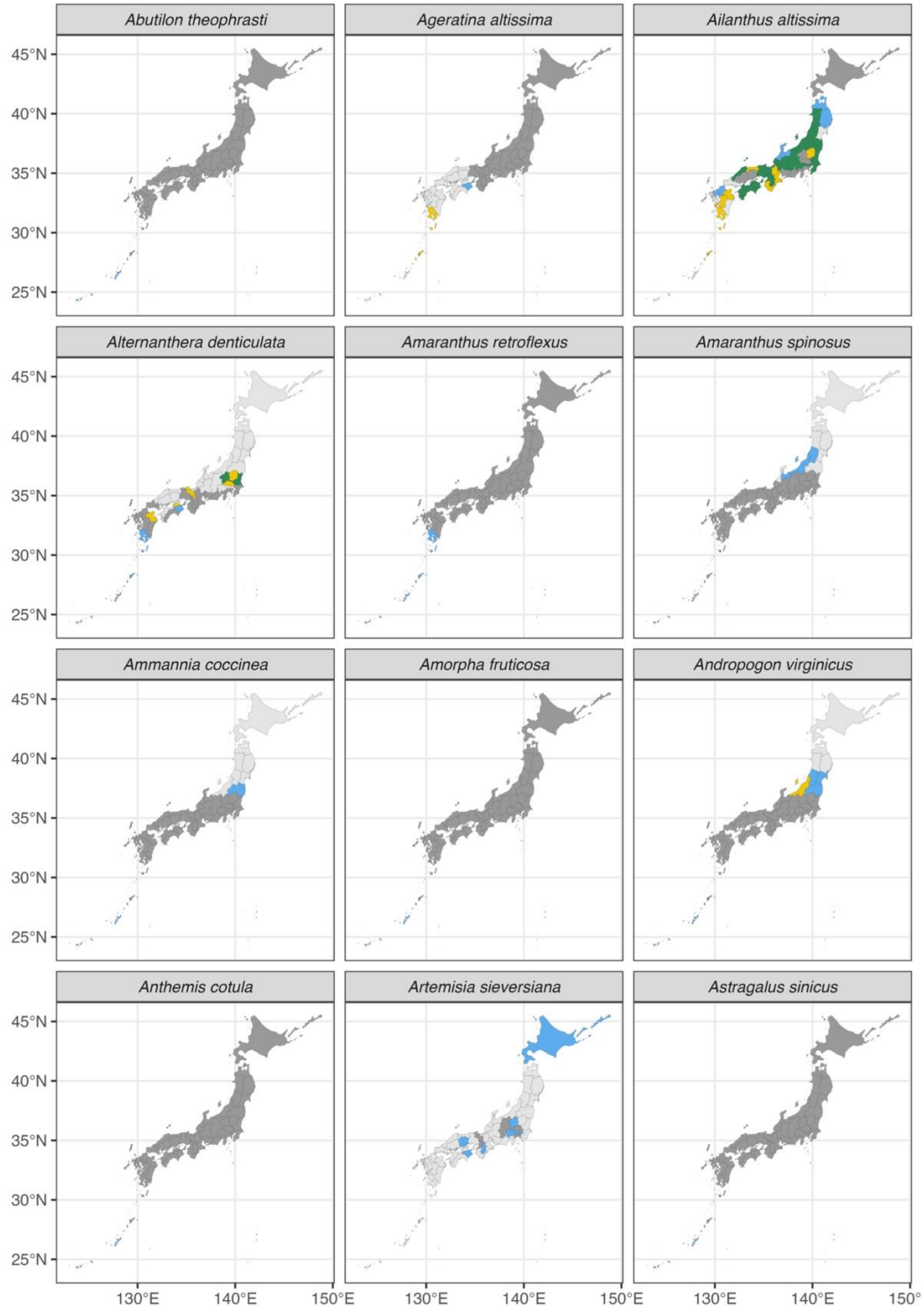
Abutilon theophrasti
Ageratina altissima
Ailanthus altissima
Alternanthera denticulata
Amaranthus retroflexus
Amaranthus spinosus
Ammannia coccinea
Amorpha fruticosa
Andropogon virginicus
Anthemis cotula
Artemisia sieversiana
Astragalus sinicus
45°N
40°N
35°N
30°N
25°N
130°E
140°E
150°E
150°

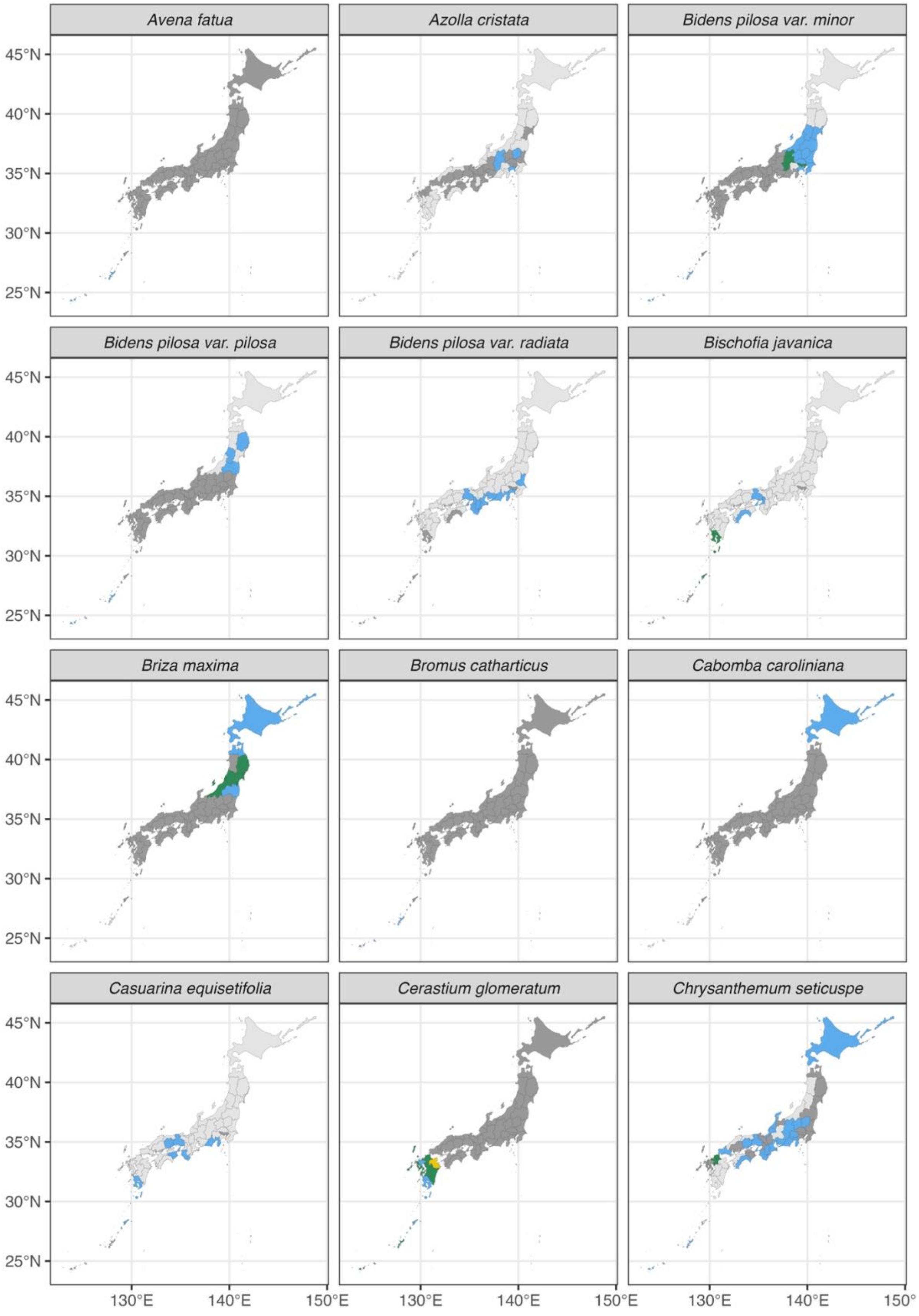
Avena fatua
Azolla cristata
Bidens pilosa var. minor
Bidens pilosa var. pilosa
Bidens pilosa var. radiata
Bischofia javanica
Briza maxima
Bromus catharticus
Cabomba caroliniana
Casuarina equisetifolia
Cerastium glomeratum
Chrysanthemum seticuspe
45°N
40°N
35°N
30°N
25°N
130°E
140°E
150°E

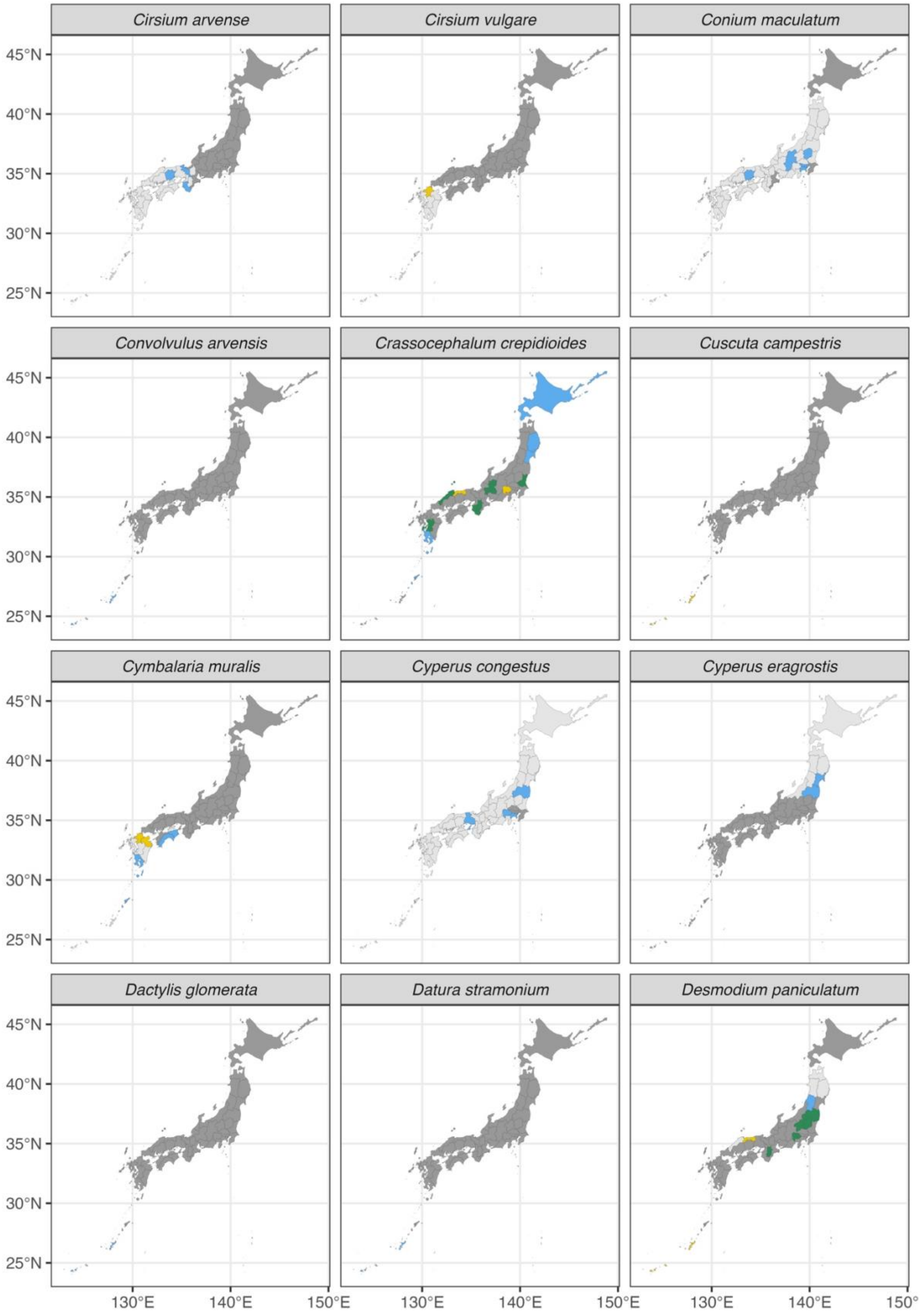
Cirsium arvense
Cirsium vulgare
Conium maculatum
Convolvulus arvensis
Crassocephalum crepidioides
Cuscuta campestris
Cymbalaria muralis
Cyperus congestus
Cyperus eragrostis
Dactylis glomerata
Datura stramonium
Desmodium paniculatum
45°N
40°N
35°N
30°N
25°N
130°E
140°E
150°E

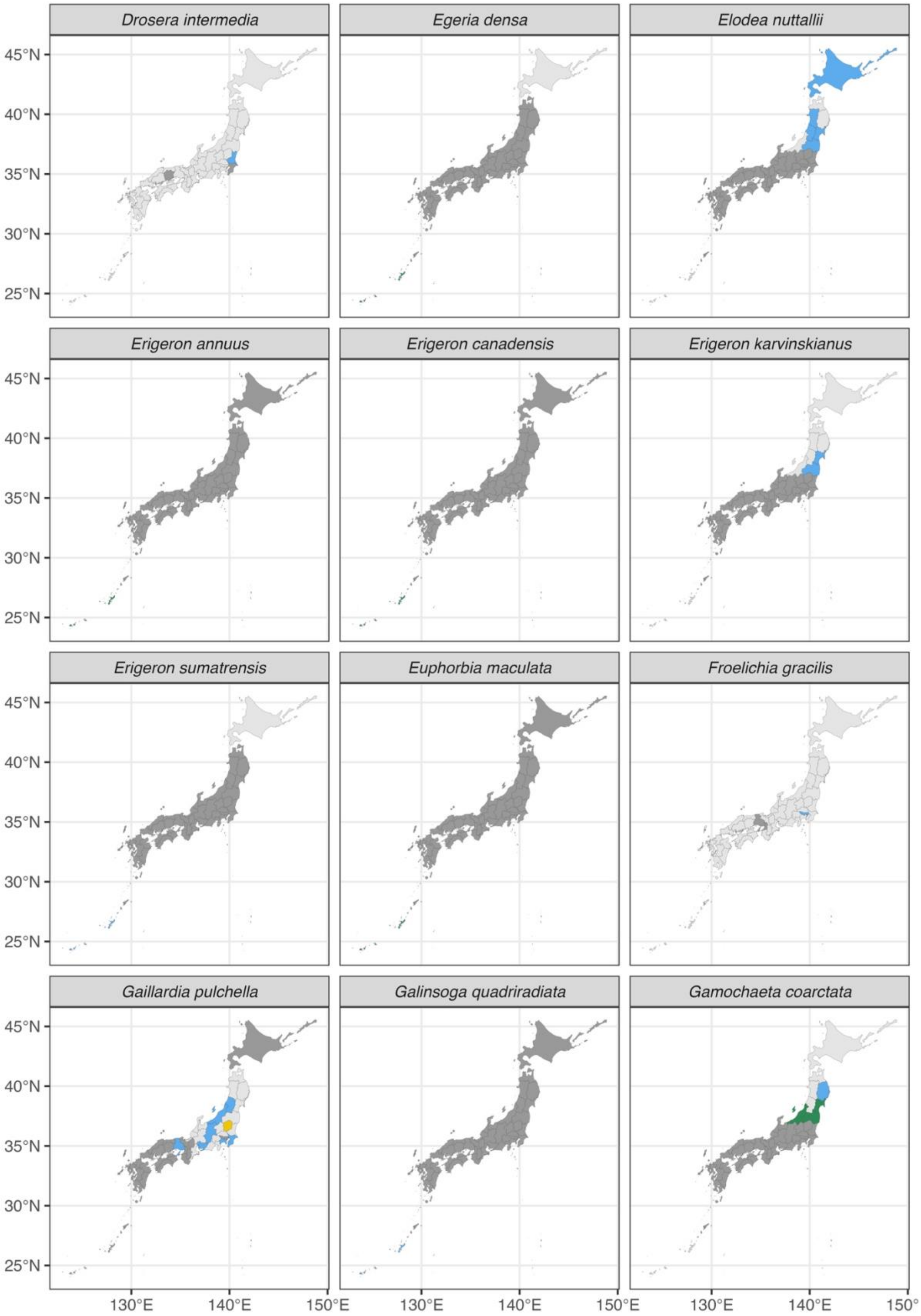

Drosera intermedia
Egeria densa
Elodea nuttallii
Erigeron annuus
Erigeron canadensis
Erigeron karvinskianus
Erigeron sumatrensis
Euphorbia maculata
Froelichia gracilis
Gaillardia pulchella
Galinsoga quadriradiata
Gamochaeta coarctata
45°N
40°N
35°N
30°N
25°N
130°E
140°E
150°E
150°

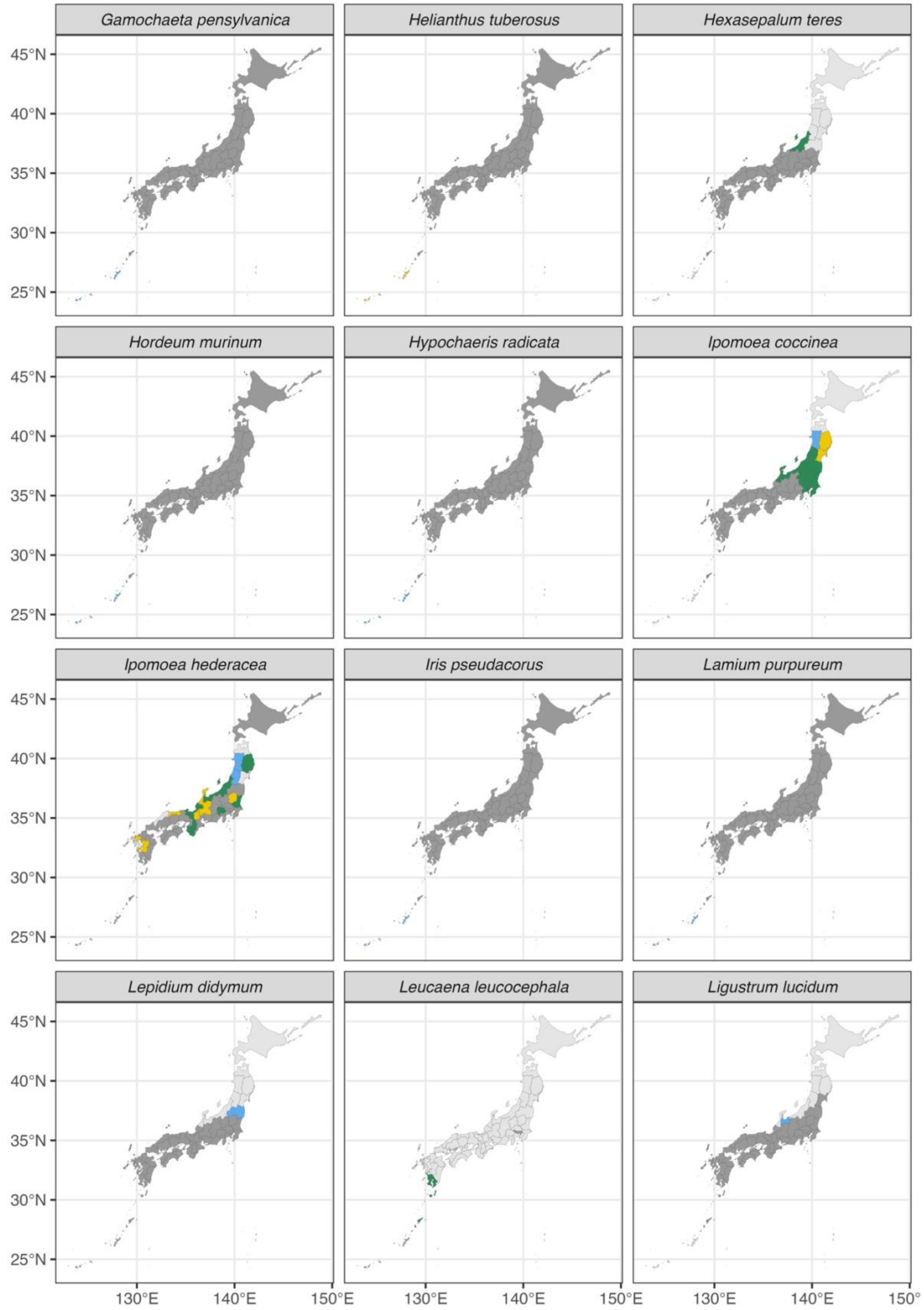
Gamochaeta pensylvanica
Helianthus tuberosus
Hexasepalum teres
Hordeum murinum
Hypochaeris radicata
Ipomoea coccinea
Ipomoea hederacea
Iris pseudacorus
Lamium purpureum
Lepidium didymum
Leucaena leucocephala
Ligustrum lucidum
45°N
40°N
35°N
30°N
25°N
130°E
140°E
150°E

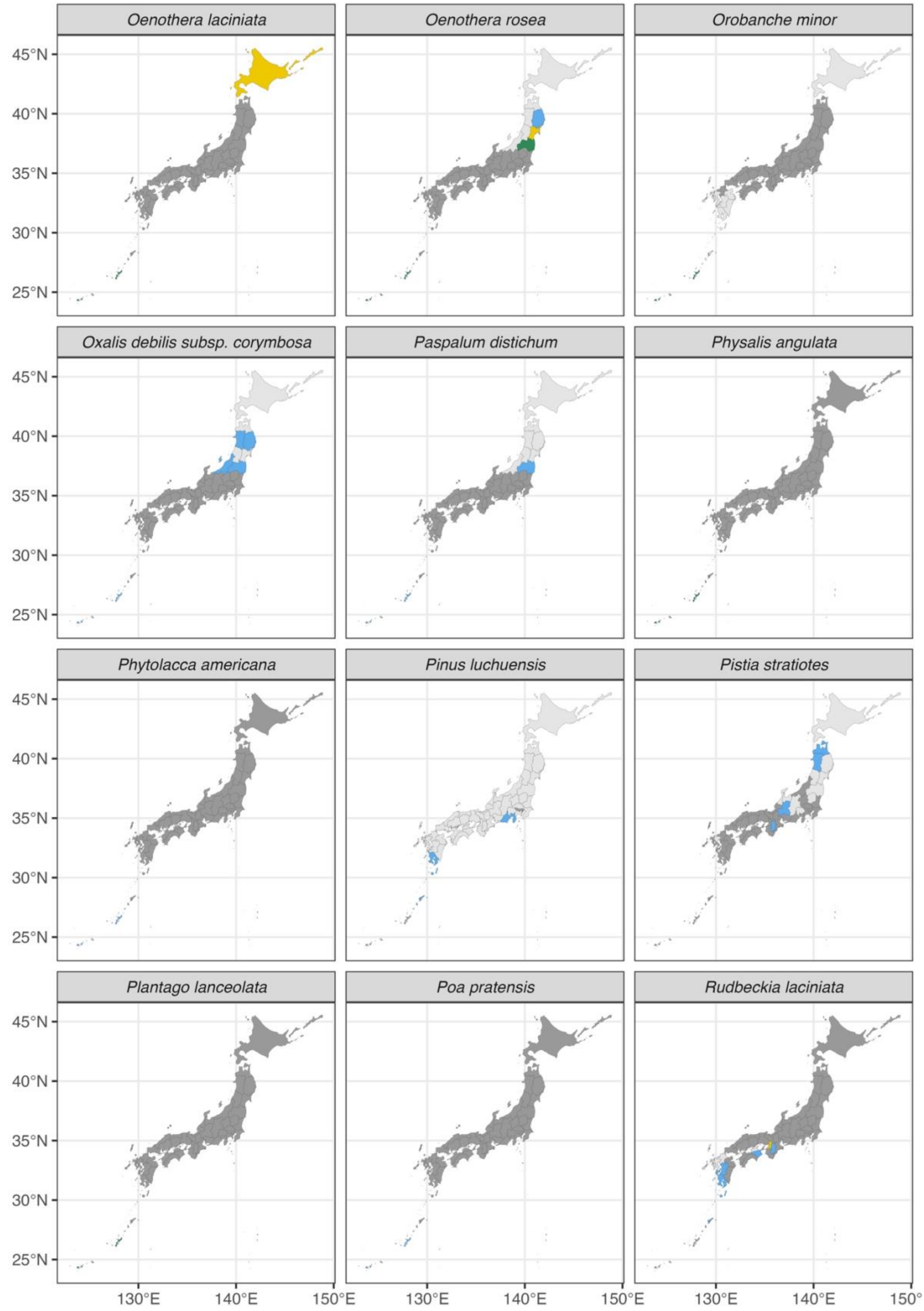

Oenothera laciniata
Oenothera rosea
Orobanche minor
Oxalis debilis subsp. corymbosa
Paspalum distichum
Physalis angulata
Phytolacca americana
Pinus luchuensis
Pistia stratiotes
Plantago lanceolata
Poa pratensis
Rudbeckia laciniata
45°N
40°N
35°N
30°N
25°N
130°E
140°E
150°E

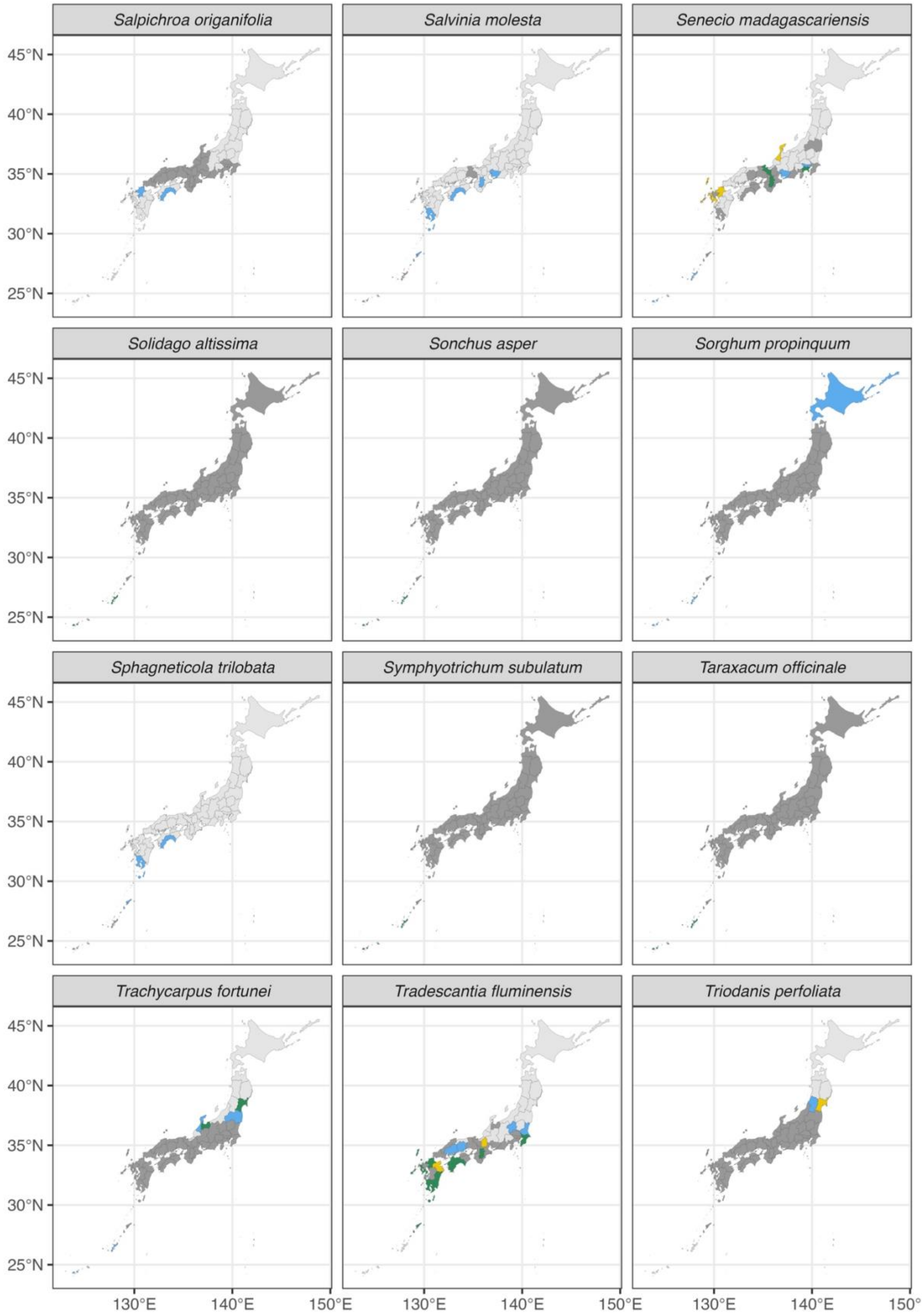
Salpichroa origanifolia
Salvinia molesta
Senecio madagascariensis
Solidago altissima
Sonchus asper
Sorghum propinquum
Sphagneticola trilobata
Symphyotrichum subulatum
Taraxacum officinale
Trachycarpus fortunei
Tradescantia fluminensis
Triodanis perfoliata
45°N
40°N
35°N
30°N
25°N
130°E
140°E
150°E
150°

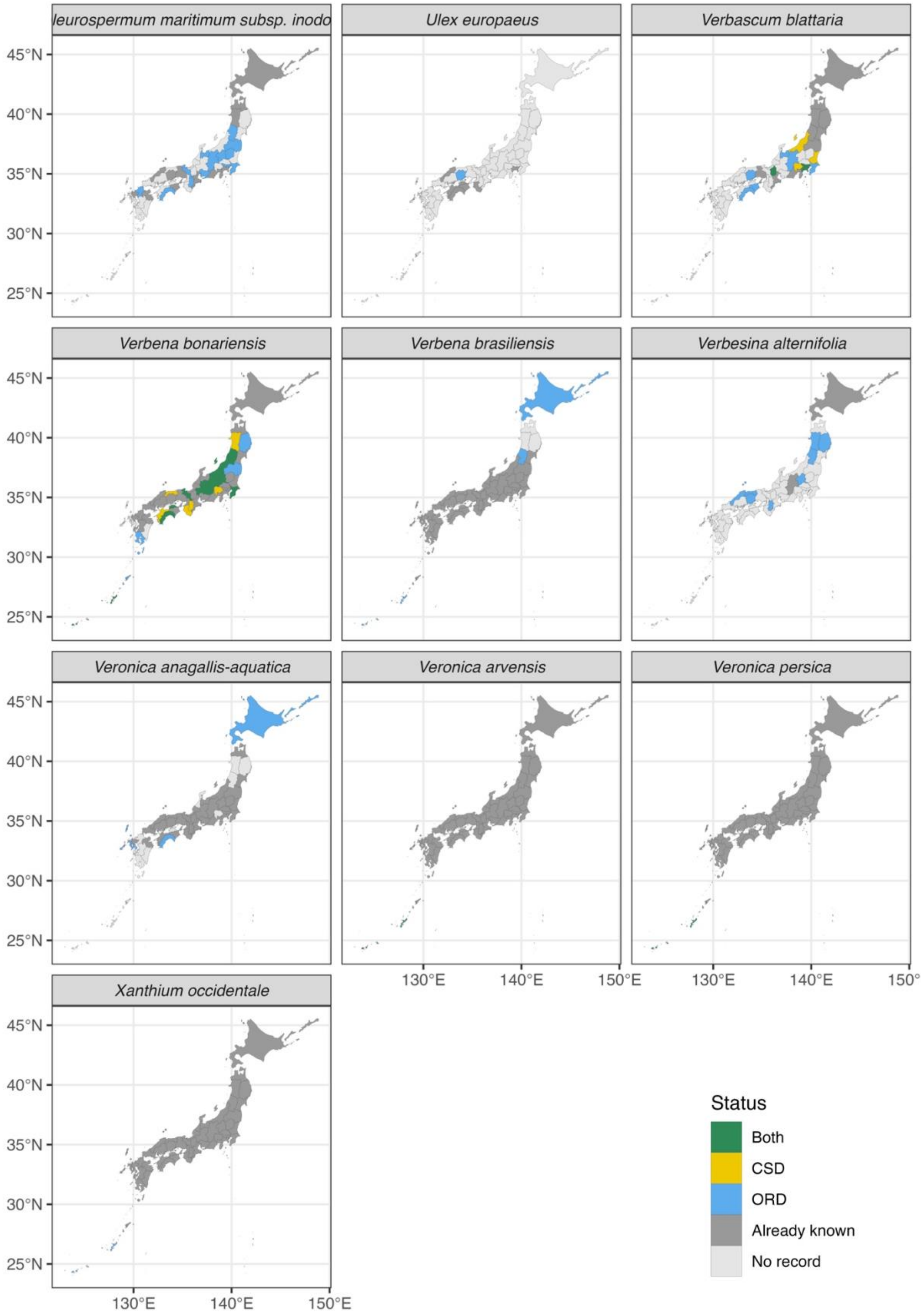

leurospermum maritimum subsp. inodo
Ulex europaeus
Verbascum blattaria
Verbena bonariensis
Verbena brasiliensis
Verbesina alternifolia
Veronica anagallis-aquatica
Veronica arvensis
Veronica persica
Xanthium occidentale
45°N
40°N
35°N
30°N
25°N
130°E
140°E
150°E
Status
Both
CSD
ORD
Already known
No record
