## Supplementary material for "Harnessing Community Science and Open Research-based Data to Track Distributions of Invasive Species in Japan": Figure S4

**Figure S4.** Updated maps of the 16 invasive insect species. The species in which we did not find new distributions were excluded. The colors of the prefectures represent the following categories: the prefecture where 1) the presence of the species had already been reported in the Invasive Species Database (ISD) (Already known), 2) the presence of species was indicated by both CSD and ORD (Both), 3) by CSD only (CSD) or 4) by ORD only, and 5) there is no data supporting the presence of the species (No record). The original shape file of the map was obtained from the Geospatial Information Authority of Japan (https://www.gsi.go.jp/top.html) and modified by the authors for analytical purposes.

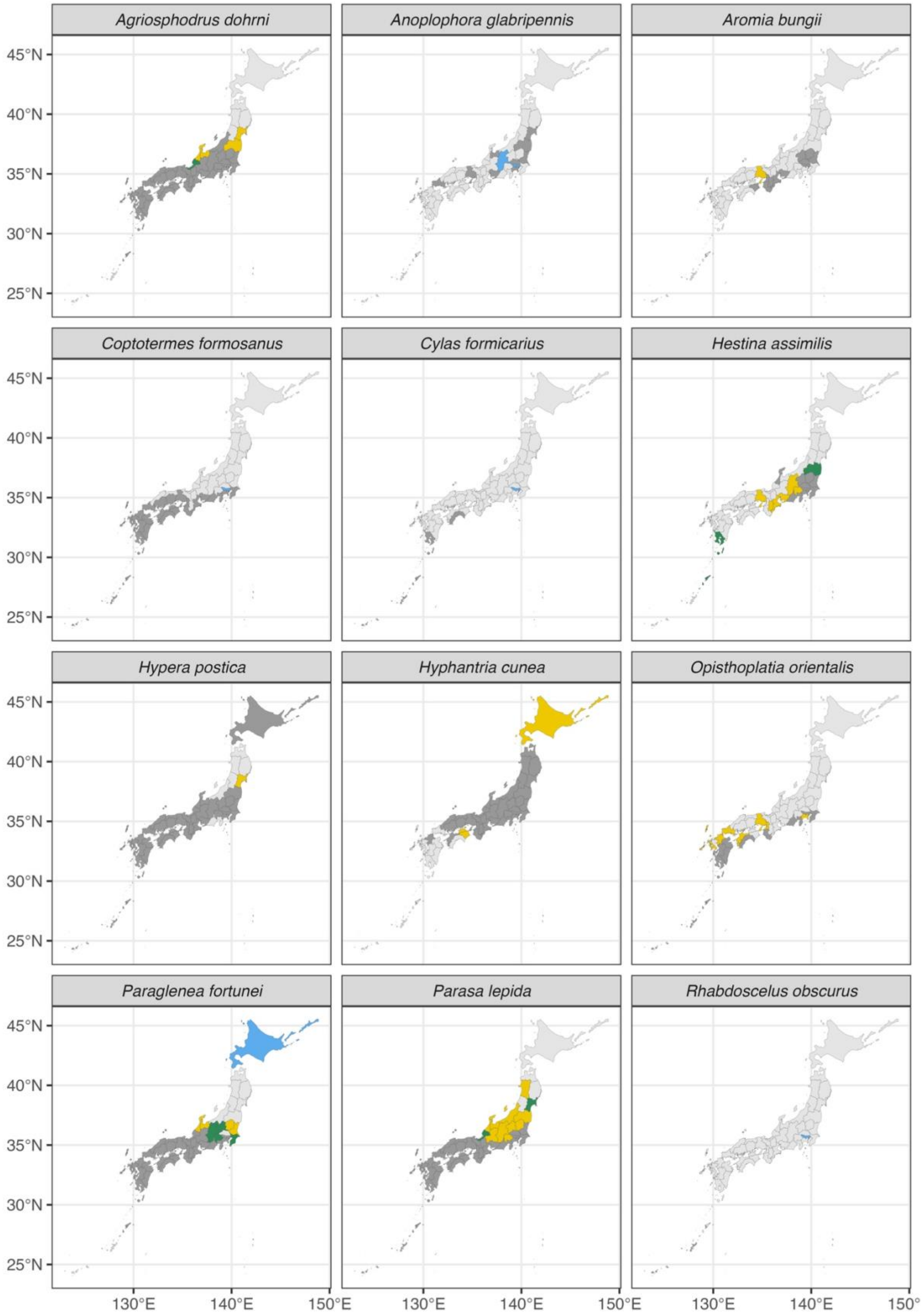
Agriosphodrus dohrni
Anoplophora glabripennis
Aromia bungii
Coptotermes formosanus
Cylas formicarius
Hestina assimilis
Hypera postica
Hyphantria cunea
Opisthoplatia orientalis
Paraglenea fortunei
Parasa lepida
Rhabdoscelus obscurus
45°N
40°N
35°N
30°N
25°N
130°E
140°E
150°E

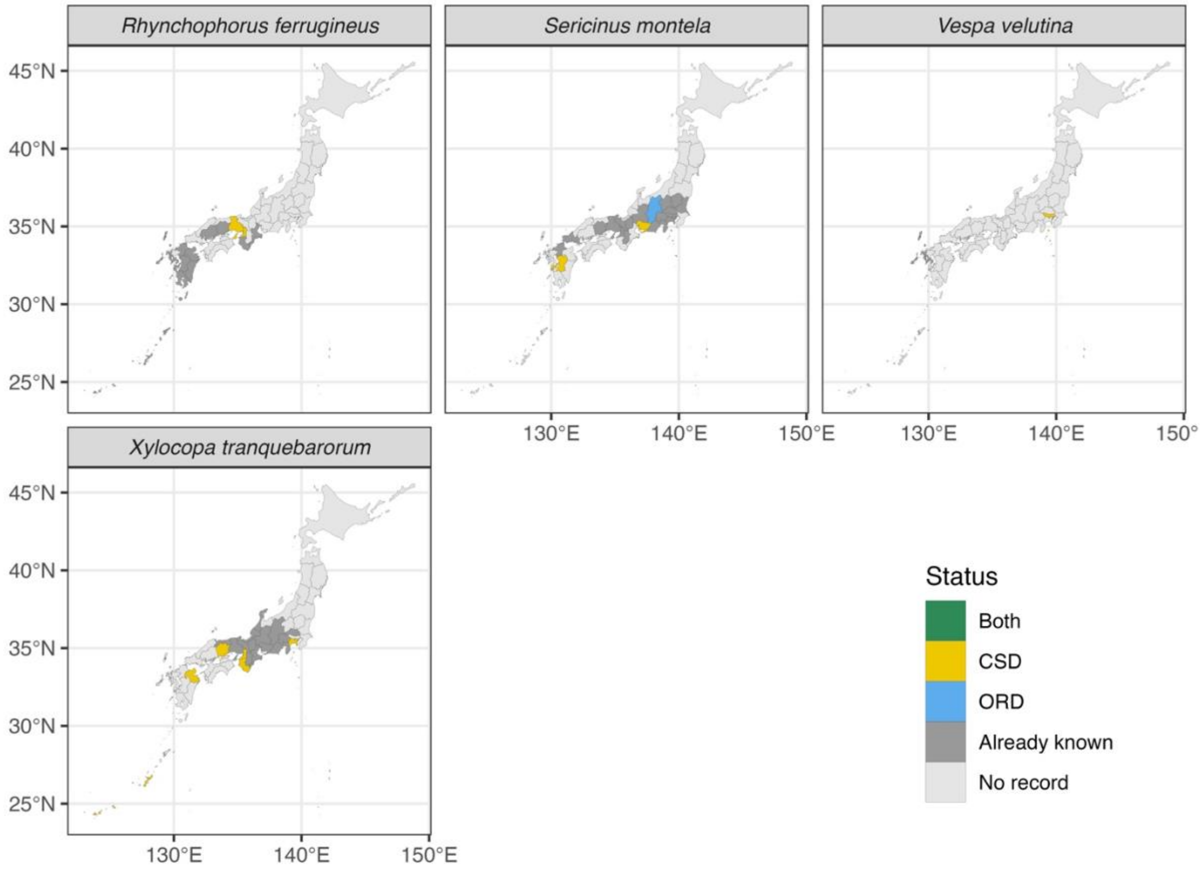
Rhynchophorus ferrugineus
Sericinus montela
Vespa velutina
Xylocopa tranquebarorum
45°N
40°N
35°N
30°N
25°N
130°E
140°E
150°E
Status
Both
CSD
ORD
Already known
No record
