## Supplementary material for "Harnessing Community Science and Open Research-based Data to Track Distributions of Invasive Species in Japan": Figure S5

**Figure S5.** Plots of the number of occurrence records and whether the occurrence of the species in the prefecture was confirmed by manual validation in Biome data for the plants (**A**) and insects (**B**). Each symbol represents a combination of species and prefecture. Three plant and insect species with high occurrences are highlighted with different colors, while others are indicated by open circles. Points are jittered to show their aggregated distribution better.

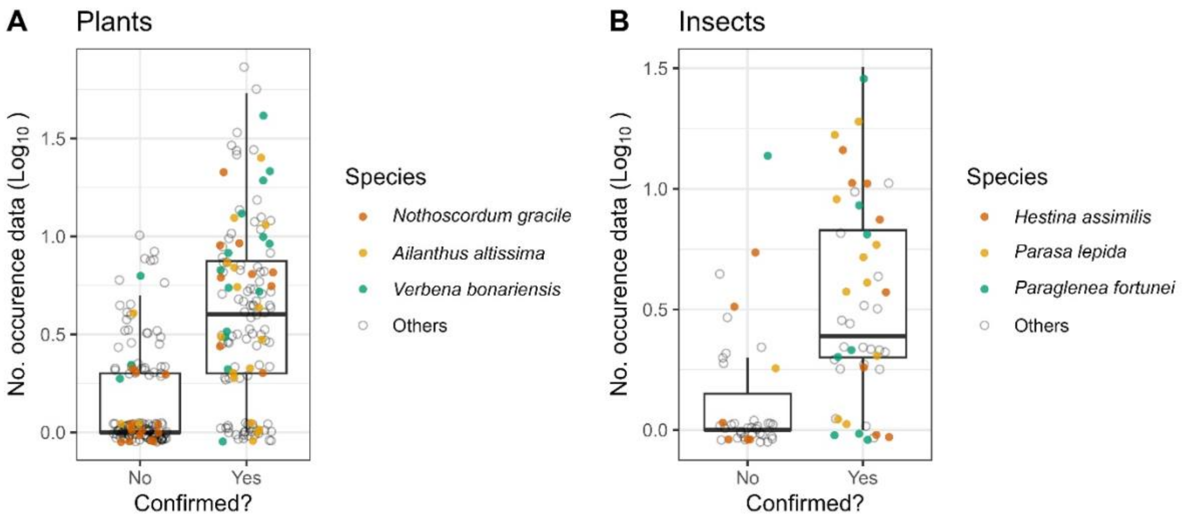
